# Development of a major histocompatibility complex class II conditional knockout mouse to study cell-specific and time-dependent adaptive immune responses in peripheral nerves

**DOI:** 10.1101/2023.07.24.550421

**Authors:** Eroboghene E. Ubogu, Jeremy A. Conner, Yimin Wang, Dinesh Yadav, Thomas L. Saunders

## Abstract

**Introduction:** Major histocompatibility complex (MHC) class II professional antigen presenting cell-naïve CD4+ T-cell receptor complex interactions are necessary for adaptive immunity. MHC class II upregulation occurs in multiple cell types in autoimmune polyneuropathy patient biopsies, necessitating precise cellular pathway studies in tissue-specific autoimmunity.

**Methods:** Cryopreserved Guillain-Barré syndrome (GBS) patient sural nerve biopsies and sciatic nerves from the severe murine experimental autoimmune neuritis (sm-EAN) GBS model were studied. Cultured conditional ready MHC Class II antigen A-alpha chain (H2-Aa) embryonic stem cells were used to generate H2-Aa^flox/+^ C57BL/6 mice. Mice were backcrossed and intercrossed to the SJL background to generate H2-Aa^flox/flox^ SJL mice, bred with hemizygous Tamoxifen-inducible von Willebrand factor Cre recombinase (vWF-iCre/+) SJL mice to generate H2-Aa^flox/flox^; vWF-iCre/+ to study microvascular endothelial cell adaptive immune responses. Sm-EAN was induced in Tamoxifen-treated H2-Aa^flox/flox^; vWF-iCre/+, H2-Aa^flox/flox^; +/+, H2-Aa^+/+^; vWF-iCre/+ and untreated H2-Aa^flox/flox^; vWF-iCre/+ adult female SJL mice. Neurobehavioral, electrophysiological and histopathological assessments were performed at predefined time points.

**Results:** Endoneurial endothelial cell MHC class II expression was observed in normal and inflamed human and mouse peripheral nerves. Tamoxifen-treated H2-Aa^flox/flox^; vWF-iCre/+ mice were resistant to sm-EAN despite extensive MHC class II expression in lymphoid and non-lymphoid tissues.

**Discussion:** A conditional MHC class II knockout mouse to study cell- and time-dependent adaptive immune responses *in vivo* is developed. Initial studies show microvascular endothelial cell MHC class II expression is necessary for peripheral nerve specific autoimmunity, as advocated by human *in vitro* adaptive immunity and *ex vivo* transplant rejection studies.

## Introduction

Primary adaptive immunity is initiated by exogenous antigen presentation to naïve CD4+ T-cells by “professional” myeloid antigen-presenting cells (APCs), e.g. dendritic cells, macrophages and B-cells, residing in secondary lymphoid organs.^1–4^ In response to antigen recognition, naïve CD4+ T-cells expand and differentiate into effector and memory T-cells. The latter persist long after initial antigen clearance, and rapidly initiate secondary tissue responses following specific antigen reappearance.^1,4,5^ Molecular mimicry is a mechanism by which infections, chemical agents and tissue injury (e.g. surgery) induce autoimmunity due to similarities between exogenous and endogenous peptides that result in autoreactive T- or B-cell activation in susceptible individuals.^3^ Patient age, sex, genetic and epigenetic factors, prior multiple pathogen exposure, vaccination status, microbiome and other environmental exposures are some molecular mimicry influencing factors.^2–4^ Dysregulated central tolerance, non-specific bystander activation, persistent antigenic stimuli and epitope shift/ drift also contribute to autoimmune disease initiation, relapse and persistence.^2^

Molecular mimicry is considered essential to GBS immunopathogenesis, with pathogenic antibodies generated against peripheral axon complex ganglioside and peripheral nerve myelin proteins/ lipids.^6–8^ Definitive pathogenic antibodies responsible for the most common GBS variant worldwide, acute inflammatory demyelinating polyradiculoneuropathy (AIDP), remain elusive. Histopathological evaluation of AIDP-variant GBS patient biopsies indicate hematogenous leukocyte trafficking with monocyte/ macrophage, CD4+ and CD8+ T-cell and B-cell/ plasma cell infiltration associated with demyelination and axonal degeneration.^9–14^ Importantly, electron-dense intercellular contacts between endoneurial endothelial cells, indicative of tight junctions, are present during pathogenic leukocyte trafficking in AIDP, supporting the notion that leukocyte trafficking across microvessels is a coordinated, sequential process (i.e. multistep paradigm of leukocyte trafficking) dependent on dynamic leukocyte-endothelial cell interactions *in vivo* as seen *in vitro*,^13,15–19^ rather passively due to a “leaky” blood-nerve barrier (BNB).

The initial directional cues for antigen-specific effector or autoreactive leukocyte trafficking into peripheral nerves during autoimmunity are unknown. How do recently activated effector leukocytes generated in secondary lymphoid organs, efficiently migrate to peripheral nerves and nerve roots alone to induce demyelination and axonal degeneration in susceptive individuals as part of aberrant adaptive immune response? Based on human *in vitro* and *ex vivo* studies, effector memory CD4+ T-cells are recruited following local microvasculature activation by inflammatory cytokines, pathogen-associated or damage-associated molecular patterns.^1,5^ Selective chemokine and cell adhesion molecules attract and facilitate leukocyte subset transmigration.^20–22^ Endothelial cell MHC Class II antigen presentation also facilitates CD4+ effector memory CD4+ T-cell activation and proliferation, with increased evidence supporting a role in antigen-specific T-cell transmigration and allograft transplant rejection.^23–27^

Increased MHC Class II (also known as human leukocyte antigen [HLA]-subtype D) expression occurs on multiple cell types: monocytes/ macrophages, Schwann cells, infiltrating T-cells, perineurial cells and BNB-forming endoneurial endothelial cells, in inflammatory neuropathy patient nerve biopsies.^28–33^ Schwann cells possess the required molecular machinery to serve as facultative APCs during adaptive immune responses *in vitro* and *in vivo*.^31,34^ A selective Schwann cell MHC class II knockout mouse model demonstrated a role for Schwann cell adaptive immune responses in preventing chronic nociception following chronic constriction nerve injury.^35^ Human and mouse BNB-forming endoneurial endothelial cells also possess the molecular machinery to serve as secondary antigen presenting cells during adaptive immune responses.^36,37^

Mouse MHC class II molecules consists of a 33 kDa α-chain covalently linked to a 28 KDa β-chain. Mice possess 2 α-chains, H2-Aa and H2-Ea, homologous to human HLA-DQA1 and HLA-DRA molecules respectively. Classical (H2-Ab1 and H2-Eb1) and nonclassical (H2-Ob [H2-Ab2] and H2-Eb2) β-chains can bind to the highly conserved α-chain to form effective antigen presentation MHC class II complexes.^38^ Interestingly, mouse strains commonly used to model human autoimmune disorders (including AIDP) such as C57BL/6, SJL and non-obese diabetic mice are H2-Ea (also known as I-E) germline knockouts (see lists from BioLegend: https://www.biolegend.com/Files/Images/media_assets/support_resource/BL_MouseAlloantigens_041116.pdf and BD Biosciences: https://www.bdbiosciences.com/content/dam/bdb/marketing-documents/mouse_alloantigens_chart.pdf). This implies that experimental autoimmunity in these mouse strains are H2-Aa-dependent. In order to determine which APCs are necessary for peripheral nerve-specific autoimmunity *in vivo*, we sought to develop a conditional H2-Aa knockout mouse strain.

## Methods

### Development of Tamoxifen-inducible microvascular-specific conditional H2-Aa knockout SJL mice

The conditional ready heterozygous H2-Aa^tm1a(KOMP)Wtsi^ embryonic stem (ES) cells, clone 1 (H2-Aa G03 (EPD0585_5_G03) derived from the JM8A3.N1 Agouti C57BL/6N mouse cell line (consisting of 61-70% euploid cells) was purchased from the University of California Davis Knockout Mouse Program (KOMP; Research Resource Identifier (RRID): MMRRC_056665-UCD; Mouse Genome Informatics: MGI:4458506). ES cell genotyping performed by KOMP demonstrated a Y chromosome, and the target was confirmed by Long Range polymerase chain reaction (PCR) at the vector’s 5’ end. Conditional potential was verified by identifying the 3’ “locus of crossing over in P1” (loxP) sequence using quantitative Taqman PCR relative to an internal genomic reference. The H2-Aa G03 ES cell line was cultured and expanded by the University of Michigan Transgenic Animal Model Core, following validated KOMP protocols (https://www.komp.org/protocols.php).

Superovulated albino C57BL/6 mouse blastocysts were microinjected with H2-Aa ES cells and transferred to pseudopregnant recipients, generating H2-Aa ES cell-mouse chimeras. All primers used in this study were produced by the UAB Heflin Center Genomics Core/ Integrated DNA technologies (www.idtdna.com). DNA amplification was performed using a Techne TC-412 Thermal cycler with specific PCR programs shown in Supplementary Table 1, unless otherwise stated. Tail snip genotyping was performed to detect mice with the complete H2-Aa transgene (i.e. the H2-Aa targeted sequence [exons 2, 3, 4 and 5’ end of exon 5] flanked by 5’ and 3’ loxP sites and neomycin resistance selection cassette) based on KOMP protocols using the following primers: 5’ loxP Forward Primer 5’-AATCTGTGCCTTGCTGATGCTTACC-3’ and Reverse Primer 5’-TCTACCATCCTGACTGCCTTGTTCC-3’ (amplicon size 652 base pairs [bp] in wildtype and 870 bp with floxed allele); 3’ loxP Forward Primer 5’-GAGATGGCGCAACGCAATTAATG-3’ and Reverse Primer 5’-CTGGCTCCAGTAAGCTCCAAAATGG-3’ (amplicon size 279 bp with floxed allele); and neomycin resistance selection cassette (amplicon size 654 bp) Forward Primer 5’-GGGATCTCATGCTGGAGTTCTTCG-3’ and Reverse Primer 5’ TCTACCATCCTGACTGCCTTGTTCC-3’.

6-week old male albino C57BL/6 H2-Aa chimeras were bred with female albino C57BL/6 flippase o (flpo) transgenic mice^39^ every 2 weeks to delete flippase recombinase target (FRT)-flanked β-galactosidase reporter and neomycin resistance selection cassette in offspring by the University of Michigan Transgenic Animal Core. This was to generate hemizygous flpo recombinase mice heterozygous for only the floxed H2-Aa targeted gene sequence (i.e. albino C57BL/6 H2-Aa^flox/+^). Flpo transgene genotyping was performed as follows: flpo Forward Primer 5’-TGAGCTTCGACATCGTGAAC-3’ and flpo Reverse Primer 5’-CAGCATCTTCTTGCTGTGG - 3’. PCR amplification was performed by adding 1.0 µL purified tail snip genomic deoxyribonucleic acid (DNA) to a mixture of 0.5 µL each of forward and reverse primers from a 10 µM stock concentration with 2.5 µL of 10X PCR buffer, 0.2 µL of 5U/ µL Taq DNA polymerase, 0.5 µL of deoxyribonucleotide triphosphate (dNTP) mix (10 mM each), 0.75 µL of 50 mM MgCl_2_ and 19.05 µL of sterile molecular grade ddH_2_O, with a total of 25 µL PCR reaction mixture.

PCR products were loaded on a 3% agarose gel in 1X TAE buffer (40 mM **T**ris base, 20 mM **A**cetic acid, and 1 mM **E**thylenediaminetetraacetic acid [EDTA]) containing 0.01% MilliporeSigma™ GelRed™ Nucleic Acid Stain (Fisher Scientific, catalog # SCT123) with electrophoresis performed at 65V for 1 hour to detect the 230 bp transgene amplicon. GeneRuler 100 bp DNA Ladder (ThermoFisher scientific, catalog # FERSM0242) was used to confirm band size for all PCR assays. Direct or reverse digital gel images were generated using an AlphaImager HP image documentation system (Cell Biosciences) attached to a Sony ICX267AL 1.39 Megapixel charged-coupled device (CCD) camera and processed using the AlphaView software program for all PCR assays.

6-week old male hemizygous flpo recombinase albino C57BL/6 H2-Aa^flox/+^ mice were bred with female albino C57BL/6 wildtype mice every 2 weeks, and genetically confirmed albino C57BL/6 H2-Aa^flox/+^ mice were backcrossed with wildtype C57BL/6 mice for 10 generations to generate congenic C57BL/6 H2-Aa^flox/+^ (officially designated as C57BL/6-*H2-Aa^tm1c(KOMP)Wtsi^*/UbeeMmmh; Mutant Mouse Resource and Research Center [MMRRC] citation ID: RRID:MMRRC_046053-MU). Genotyping was performed to detect the H2-Aa targeted gene sequence at the 5’ and 3’ loxP sites using the previously described KOMP protocol.

In order to study adaptive immune responses in severe murine experimental autoimmune neuritis (sm-EAN), C57BL/6 H2-Aa^flox/+^ mice were backcrossed to the SJL background by speed congenics, performed at Charles River using their proprietary Marker-Assisted Accelerated Backcrossing (MAX-BAX^®^) protocol. Due to H2-Aa sequence differences in exon 2 between SJL and C57BL/6 mice, H2-Aa SJL transgene genotyping was performed as follows: 5’ loxP Forward Primer 5’-GCACCATGAATCTGCAAGTGAACA-3’ and Reverse Primer 5’-TCTACCATCCTGACTGCCTTGTTCC-3’ (amplicon size: 652 bp wildtype, and 870 bp floxed allele); 3’ loxP Forward Primer 5’-GAGATGGCGCAACGCAATTAATG-3’ and Reverse Primer 5’-CTGGCTCCAGTAAGCTCCAAAATGG-3’ (amplicon size 279 bp floxed allele, unchanged from C57BL/6). PCR amplification was performed by adding 1.0 µL purified tail snip genomic DNA to a mixture of 0.5 µL each of forward and reverse primers from a 20 µM stock concentration with 12.5 µL of 2X Promega GoTaq^®^ Green Master Mix (catalog # PRM5123) and 10.5 µL sterile ddH_2_O, with a total of 25 µL PCR reaction mixture. PCR products were loaded on a 2% agarose gel in 1X TAE buffer containing 0.01% MilliporeSigma™ GelRed™ Nucleic Acid Stain (Fisher Scientific, catalog # SCT123) with electrophoresis performed at 200V for 30 minutes for the 3’ loxP sequence reaction and 40 minutes of the 5’ loxP sequence reaction. Gels were visualized, imaged and processed as previously described.

Following complete backcross, SJL H2-Aa^flox/+^ mice were intercrossed to generate SJL H2-Aa^flox/flox^ mice (SJL.B6-*H2-Aa^tm1c(KOMP)Wtsi^*/UbeeMmmh; RRID: MMRRC_071292-MU). In order to study potential endothelial cell adaptive immune responses in sm-EAN, SJL H2-Aa^flox/flox^ mice were bred with hemizygous Tamoxifen-inducible von Willebrand factor Cre recombinase SJL mice (vWF-iCre/+; SJL.Cg-Tg(VWF-Cre/ERT2)C1014/UbeeMmmh; RRID: MMRRC_071293-MU), in which Cre recombinase expression is driven by a human microvascular-specific von Willebrand factor (vWF) promoter DNA sequence^40,41^ after Tamoxifen injection. Sciatic nerve microvascular endothelial cell Cre expression was verified using two fluorescent reporter mouse strains^42–44^ (Supplementary Figure 1). Detailed online information describing this mouse strain is available (https://doi.org/10.1101/2023.07.24.550419). Cre recombinase-mediated microvascular endothelial cell H2-Aa gene deletion occurs in SJL H2-Aa^flox/flox^; vWF-iCre/+ mice, i.e. a conditional microvascular endothelial cell-specific MHC Class II knockout mouse model.

Genotyping to detect the vWF-iCre transgene was performed as follows: vWF-iCre Forward Primer 5’-AATCTTTTCTCCTGCTTTAAAGAAATGTT-3’ and vWF-iCre Reverse Primer 5’-ATCTGTGACAGTTCTCCATCAGGGATCT-3’ (amplicon size 604 bp). PCR amplification was performed by adding 1.0 µL purified tail snip genomic DNA to a mixture of 0.5 µL each of forward and reverse primers from a 10 µM stock concentration with 12.5 µL of 2X Promega GoTaq^®^ Green Master Mix (catalog # PRM5123) and 10.5 µL of sterile ddH_2_O, with a total of 25 µL PCR reaction mixture.

### Development of microvascular endothelial cell-specific MHC class II conditional knockout SJL mice

The strategy used to generate H2-Aa^flox/flox^; vWF-iCre/+ SJL mice, and genotyping data showing albino C57BL/6 H2-Aa ES cell chimera mice with the target construct; hemizygous flpo recombinase albino C57BL/6 H2-Aa^flox/+^ mice with excision of the FRT-flanked β-galactosidase reporter and neomycin resistance selection cassette; derived C57BL/6 H2-Aa^flox/+^ mice during backcross to congenic C57BL/6 background, followed by backcross and intercross to generate SJL H2-Aa^flox/flox^ mice that were mated with Tamoxifen-inducible hemizygous vWF-iCre/+ SJL mice to generate H2-Aa^flox/flox^; vWF-iCre/+ SJL mice, are shown (**Figure 1**).

**Figure 1.**
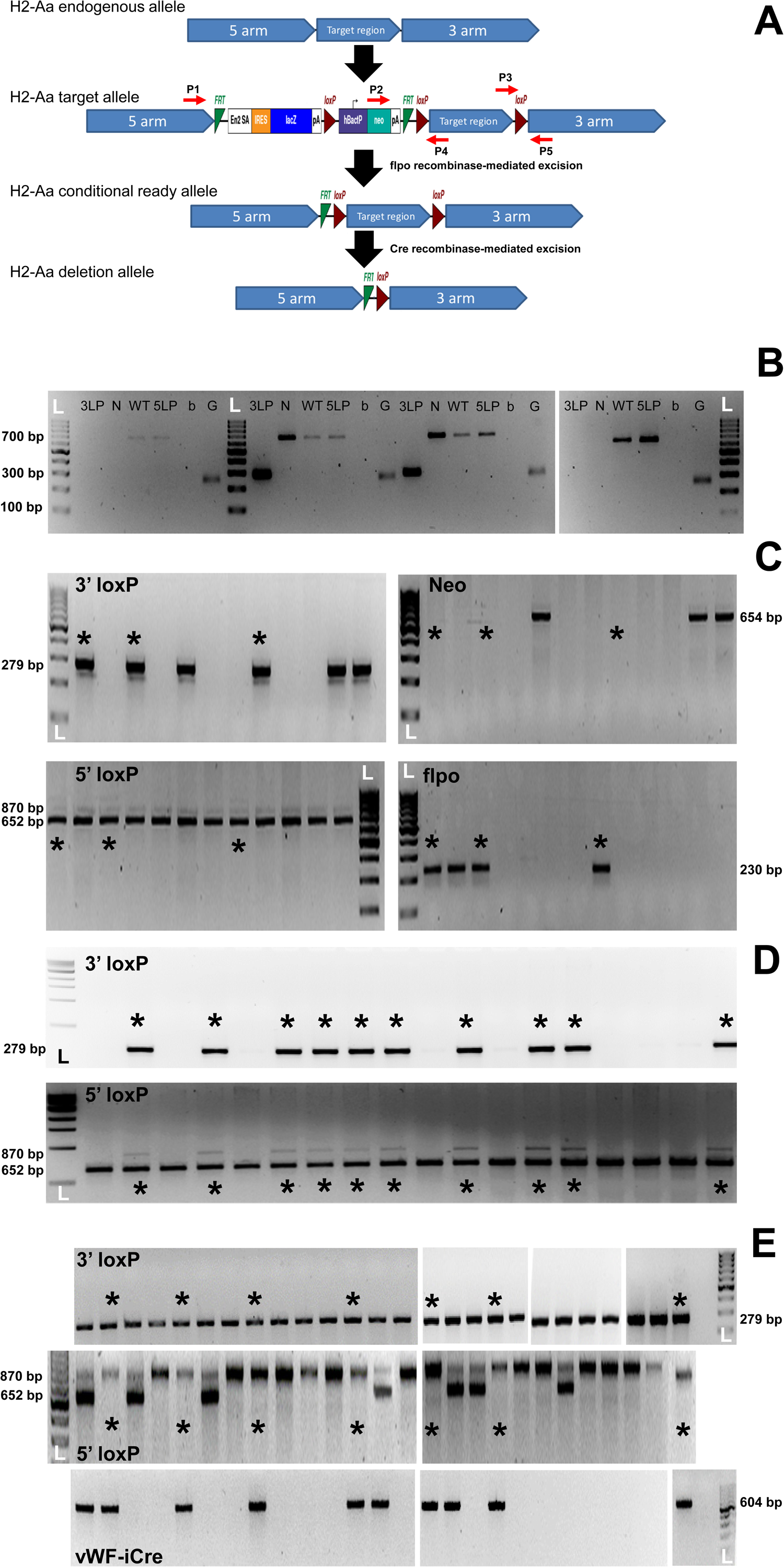
Development of microvascular endothelial-cell specific MHC class II conditional knockout SJL mouse strain. The strategy used to conditionally knockout MHC class II (H2-Aa) based on the KOMP-designed DNA construct is shown in **A**. The H2-Aa targeted allele was constructed by placing loxP sequences on the 5’ and 3’ ends of gene target region, and FRT-flanked sequences between β-galactosidase reporter and neomycin resistance selection cassette inserted 5’ to the H2-Aa targeted DNA. Following flpo recombinase-mediated excision, the conditional ready allele is produced. Excision at the loxP sites generates the H2-Aa knockout allele in cells/ tissues induced to express Cre recombinase. Representative digital reverse gel images from H2-Aa ES cell chimera albino C57BL/6 mouse tail snip genotyping for the 3’ loxP (3LP) sequence, neomycin selection cassette (N or Neo), wildtype allele (WT) and 5’ loxP (5LP) sequence, with blank (b) and glyceraldehyde-3-phosphate dehydrogenase (G) internal controls is shown in **B**. Mice with the intact targeted H2-Aa construct show PCR products at 3LP, N, WT, 5LP and G. Representative digital reverse gel images from offspring of male albino C57BL/6 H2-Aa ^flox/+^ chimera mice bred with female albino C57BL/6 flpo transgenic mice genotyped for the H2-Aa 3’ loxP, Neo and 5’ loxP sequences, and the flpo transgene show albino C57BL/6 flpo recombinase H2-Aa^flox/+^ mice with excision of the β-galactosidase reporter and neomycin resistance selection cassette, are shown in **C** (asterisk). Following mating with albino C57BL/6 mice to select heterozygous mice without the flpo transgene, albino C57BL/6 H2-Aa^flox/+^ mice were backcrossed to the C57BL/6 background, to generate congenic C57BL/6 H2-Aa^flox/+^ mice, as shown in **D** (asterisk). Following backcross to the congenic SJL background and intercross to generate SJL H2-Aa^flox/flox^ mice; these mice were bred with hemizygous Tamoxifen-inducible vWF-iCre transgenic mice. Representative gel images following tail snip genotyping of the SJL H2-Aa 3’ loxP and 5’ loxP sequences and the vWF-iCre transgene show SJL H2-Aa^flox/flox^; vWF-iCre/+ mice (asterisk) in **E**. L represents DNA size ladder and numbers indicate size markers or size of expected PCR products in B-E.

### Severe murine experimental autoimmune neuritis

8-9 week old female SJL H2-Aa^flox/flox^; vWF-iCre/+ mice were injected with 100 mg/kg Tamoxifen in corn oil i.p. for 5 consecutive days (resulting in expected maximal Cre-mediated DNA recombination 21 days after Tamoxifen injection, as demonstrated in mouse astroglia)^45^ to delete H2-Aa in microvascular endothelial cells. Similarly Tamoxifen-injected age-matched female SJL H2-Aa^flox/flox^; +/+ mice (that lack the hemizygous vWF-iCre transgene), and H2-Aa^+/+^; vWF-iCre/+ (that lack H2-Aa floxed alleles but possess the vWF-iCre transgene), as well as corn oil only-treated H2-Aa^flox/flox^; vWF-iCre/+ mice served as controls. 24 days after the final Tamoxifen injections, sm-EAN was induced via subcutaneous injections of bovine peripheral nerve myelin (BPNM) emulsified in Complete Freund Adjuvant (CFA), with Pertussis toxin and recombinant mouse interleukin (IL)-12 serving as co-adjuvants, as previously published.^13,46–49^ 100% disease induction and progression occurs reliably in female mice. Daily weights and neurobehavioral tests (neuromuscular severity score, NMSS) were performed in experimental mice for 28 days (to expected peak severity with plateau), as previously published.^13,47–49^

### Mouse motor nerve electrophysiology and tissue harvest

Bilateral sciatic and dorsal caudal tail (DCT) motor nerve electrophysiology was performed on day 28 post-induction under 100 mg/kg ketamine/ 10 mg/kg xylazine i.p, anesthesia using a portable electrodiagnostic system (Keypoint v5.11; Alpine Biomed Corp., Fountain Valley, California, USA), as previously published.^13,47,49,50^ Immediately afterwards, mice were euthanized under deep anesthesia via cervical disarticulation, and the sciatic nerves, proximal tail, inguinal lymph nodes and spleen were harvested from each mouse. The harvested tissues were immediately embedded in Scigen Tissue Plus^TM^ Optimum Cutting Temperature^®^ (OCT) Compound (Fisher Scientific, catalog # 23-730-571) or placed in cryovials and stored at −80°C until processed, or placed in fixative (3% glutaraldehyde [GTA] in 0.1M phosphate buffer or 4% paraformaldehyde [PFA] in 1X phosphate buffered saline [PBS]) in glass vials at room temperature (RT) for further processing.

### Sciatic nerve morphological assessment

A portion of the left sciatic nerve from each mouse was fixed for 12-15 hours in 3% GTA in 0.1M phosphate buffer at RT, post-fixed in 1% osmium tetroxide, and embedded in epoxy resin (plastic). 0.75 µm sections from 3-4 regions separated by ≥ 1000 µm were generated, mounted on glass slides, and stained with methylene blue to evaluate sciatic nerve morphology (axonal loss, demyelination, inflammation), as previously published.^13,47,49,51^ Slides were visualized and representative digital photomicrographs taken using a Zeiss Axio Lab.A1 upright light microscope attached to a Nikon DS-Fi2 digital camera and processed using the Nikon Elements software.

### Proximal tail decalcification and antigen retrieval

Proximal tail sections were fixed in 4% PFA in 1X PBS for 24 hours, rinsed three times for 10 minutes in 1X PBS and decalcified in 10% EDTA in 2% PFA in 1X PBS at 37°C for 2 weeks with continuous shaking at 43 revolutions per minute in a Bellco SciERA hot orbital shaker (catalog # 7745-22110), with decalcification buffer changed every 24 hours.^52^ Decalcified tail sections were washed with 1X PBS, embedded in OCT compound and stored at −80°C until processed. 50 µm thick axial sections were placed on Fisherbrand™ ColorFrost™ Plus slides (Fisher Scientific, catalog # 12-550-18), air dried for 12-15 hours overnight at RT, and stored for ∼6 hours at - 20°C. Following air drying at RT for 1 hour, antigen retrieval was performed by incubating sections with 1% Sodium Dodecyl Sulfate (SDS) in 1X PBS for 5 minutes at RT.^53^ Indirect immunohistochemistry was performed to detect DCT nerve and cutaneous inflammation.

### Indirect fluorescent immunohistochemistry

OCT-cryopreserved archived sural nerve biopsies from two adults (aged 63 and 70) with histological evidence of an acute demyelinating neuritis consistent with AIDP-variant GBS, and two age- and sex-matched histologically normal controls were obtained from the Shin J. Oh Muscle and Nerve Histopathology Laboratory, UAB. OCT-cryopreserved sciatic nerves from a normal adult female SJL mouse and sm-EAN-affected mice from disease onset (day 10 post-induction) and during the early effector phase (day 15 post-induction) from prior experiments^47^ were also obtained. These nerves were studied to confirm microvascular endothelial cell MHC Class II expression in normal and inflamed human and mouse peripheral nerves. Leukocyte MHC Class II expression in the sciatic nerves, tails, inguinal lymph nodes and spleens of Tamoxifen-treated female SJL H2-Aa^flox/flox^; vWF-iCre/+ mice (with microvascular endothelial cell H2-Aa gene deletion) and SJL H2-Aa^flox/flox^; +/+ mice (representative of mice with unaltered microvascular endothelial cell H2-Aa gene expression) following sm-EAN induction was studied to evaluate for peripheral nerve inflammation and retention of MHC class II expression in lymphoid and nonlymphoid organs.

Antibodies and lectin used for immunohistochemistry are listed in Supplementary Table 2. 10 µm thick axial sections were mounted on Fisherbrand™ Superfrost™ or ColorFrost™ Plus slides (Fisher Scientific, catalog # 12-550-14G and 12-550-18), air dried for 20-60 minutes at RT, fixed with ice cold acetone at −20°C for 10 minutes and washed twice with 1X Dulbecco’s PBS (D-PBS) for 10 minutes. Sections were blocked with 10% normal goat serum (NGS) in 1X D-PBS, followed by incubation with primary antibodies in 2% NGS in 1X D-PBS at RT for 1 hour. After washing with 2% NGS in 1X D-PBS for 10 minutes, sections were incubated with secondary antibodies/ fluorescent lectin in 2% NGS in 1X D-PBS for one hour at RT in the dark. After washing with 1X D-PBS for 10 minutes, sections were stained with 0.45 µM 4′,6-diamidino-2-phenylindole (DAPI) in 1X PBS for 5 minutes at RT in the dark to detect nuclei, and washed once with 1X D-PBS for 5 minutes. 50 µm thick fixed and decalcified axial cryostat tail sections were processed as above after antigen retrieval and 10 minute air drying, with 1X PBS as diluent and wash buffer. Sections were mounted using ProLong^TM^ Gold Antifade aqueous mounting medium (ThermoFisher Scientific, catalog # P36930) or VECTASHIELD^®^ antifade mounting medium (Vector Laboratories catalog # H100010) and coverslips placed and sealed with nail polish. Slides from the same experiments were processed immediately or stored at 4°C overnight for processing the following day.

Slides were visualized using a Nikon ECLIPSE Ci upright epifluorescent microscope attached to a Nikon DS-Qi2 monochromatic camera. Digital photomicrographs were generated and images were merged using the Nikon NIS Elements AS software program, as previously published.^13,51,54^

### Statistical Analysis

Investigator-blinded data analyses were performed for all parameters using the GraphPad Prism^®^ 6 statistical program (GraphPad Software, Inc., La Jolla, CA). The Mann-Whitney U-test or the Wilcoxon-Kruskall’s Rank Sum Test was used to determine statistically significant differences between non-parametric variables while the one- or two-tailed unpaired Student’s/Welch’s t-test (or one-way analysis of variance) was used to evaluate parametric variables based on the Shapiro-Wilk test of normality (including measures of skew and kurtosis). The Holm-Sidak’s test or Bonferroni post-test correction was used for multiple parametric variable comparisons, as previously published. Means are displayed, with individual data points or variations of the mean depicted as standard errors shown. Statistical significance is defined as a p-value < 0.05.

## Results

### MHC Class II expression in normal and inflamed human and mouse peripheral nerves

HLA-DR was expressed by BNB-forming endoneurial endothelial cells and cells with profiles consistent with endoneurial macrophages and innerlayer perineurial myofibroblasts in histologically normal adult sural nerves (**Figure 2 and Supplementary Figure 2**). Increased HLA-DR expression was observed in AIDP, with diffuse expression by cells with profiles consistent with Schwann cells, infiltrating leukocytes, endoneurial and epineurial vascular endothelial cells and perineurial myofibroblasts (**Figure 2 and Supplementary Figure 2**), as previously published.^28–33^ H2-A was expressed by BNB-forming endoneurial endothelial cells and rare cells with profiles consistent with endoneurial macrophages in normal female SJL mouse sciatic nerves. There was increased expression by endoneurial endothelial cells at sm-EAN disease onset that persisted to the effector phase, associated with infiltrating leukocytes and perineurium expression (**Figure 2 and Supplementary Figure 2**). These studies show that MHC class II is constitutively expressed by BNB-forming endothelial cells in human and mouse peripheral nerves and upregulated during acute demyelinating neuritis in association with other peripheral nerve cells.

**Figure 2.**
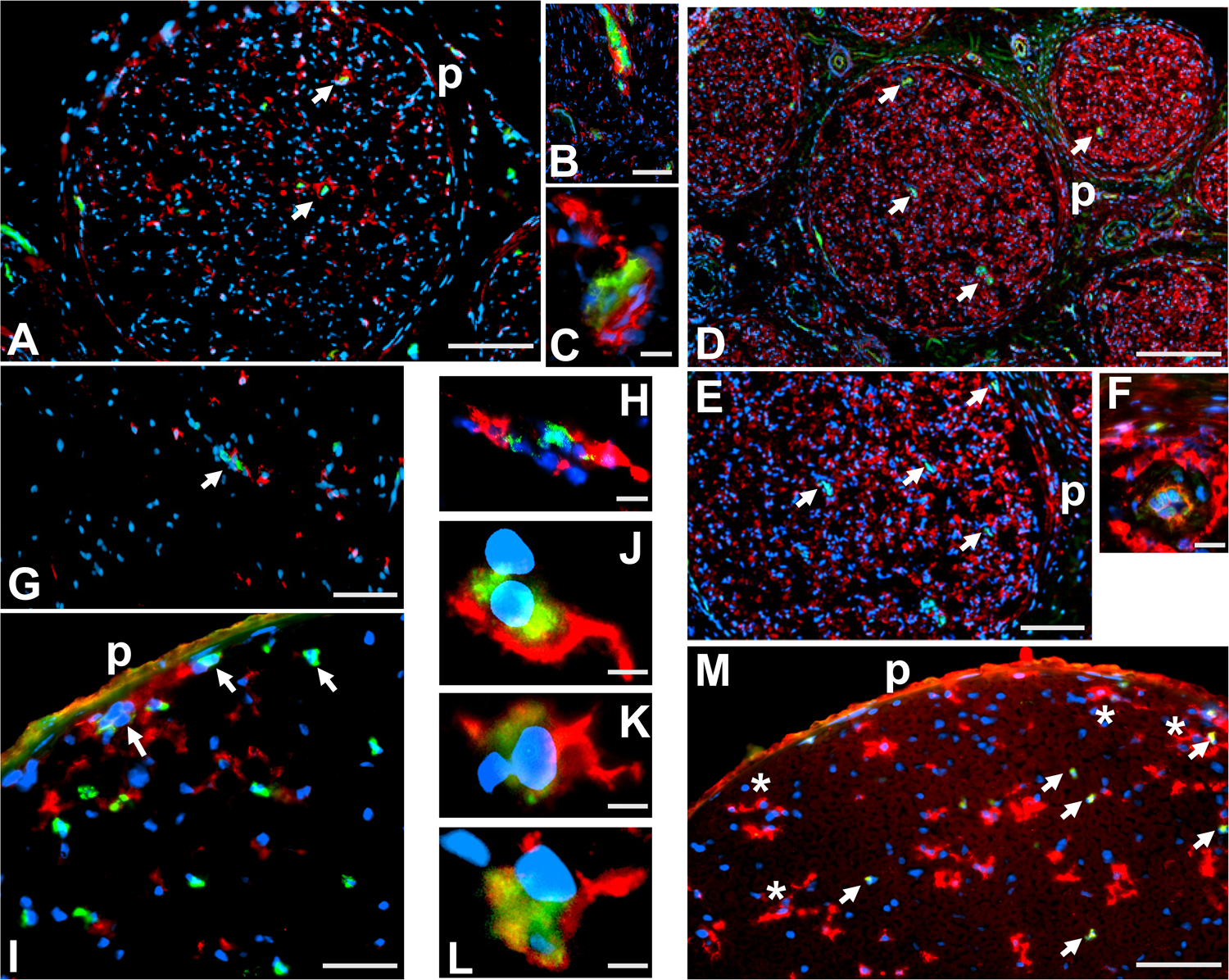
MHC Class II expression in normal and inflamed human and mouse peripheral nerves. Representative merged digital photomicrographs show HLA-DR expression (red fluorescence) by BNB-forming endoneurial endothelial cells (green fluorescence) in a histologically normal adult human sural nerve at low magnification (**A**, white arrows), with expression in the inner perineurium (p) also seen. Higher magnification images show normal endoneurial endothelial cell membrane and cytoplasmic HLA-DR expression (**B-C**). Increased diffuse HLA-DR expression (red fluorescence) is seen in AIDP patient sural nerves involving endoneurial endothelial cells (green fluorescence)cells with profiles consistent with Schwann cells and infiltrating leukocytes, as well as (**D-E**, white arrows) and the perineurium (p). Higher magnification images show endoneurial microvessel luminal (endothelial cell) and basement membrane HLA-DR expression in AIDP (**F**). H2-A expression (red fluorescence) by BNB-forming endoneurial endothelial cells (green fluorescence) is also seen in an adult female SJL mouse sciatic nerve (**G**, white arrow) and cells with profiles consistent with endoneurial macrophages, with a higher magnification image showing normal endoneurial endothelial cell luminal membrane and cytoplasmic H2-A expression (**H**). Increased H2-A expression (red fluorescence) is seen at sm-EAN onset, with expression by endoneurial endothelial cells (green fluorescence; white arrows), perineurium (p) and cells consistent with infiltrating leukocytes (**I**). Higher magnification images at sm-EAN onset show infiltrating cells associated with endoneurial microvessel H2-A expression (**J-L**), suggesting a trafficking role for H2-A at sm-EAN onset. H2-A expression (red fluorescence) by endoneurial endothelial cells (green fluorescence; white arrows), the perineurium (p) and profiles consistent with infiltrating leukocytes (asterisk) with increased diffuse endoneurial H2-A background expression is observed during the early sm-EAN effector phase (**M**). Scale bars: A = 100 μm, B-C = 20 μm, D = 200 μm, E = 100 μm, F = 5 μm, G = 50 μm, H = 10 μm, I = 25 μm, J-L = 5 μm, M = 50 μm.

### Effect of microvascular endothelial cell-specific MHC Class II conditional knockout on sm-EAN

The experimental design to study sm-EAN in age-matched female Tamoxifen-treated SJL H2-Aa^flox/flox^; vWF-iCre/+, SJL H2-Aa^flox/flox^; +/+ and H2-Aa^+/+^; vWF-iCre/+ and corn oil only-treated H2-Aa^flox/flox^; vWF-iCre/+ mice is shown in **Figure 3A**. H2-Aa^flox/flox^; vWF-iCre/+ mice were resistant to sm-EAN based on NMSS scores, with tail weakness being the only abnormality seen in 3 of 8 mice studied. All other experimental mice with unaltered microvascular endothelial cell H2-Aa gene expression fully developed sm-EAN (**Figure 3B**). Although Tamoxifen-treated female SJL H2-Aa^flox/flox^; vWF-iCre/+ mice appear to weigh less than mice with unaltered H2-Aa gene expression (**Figure 3C**), no significant difference in mean % weight change was observed between sm-EAN experimental mice (**Figure 3D**), consistent with prior reports.^47,49^

**Figure 3.**
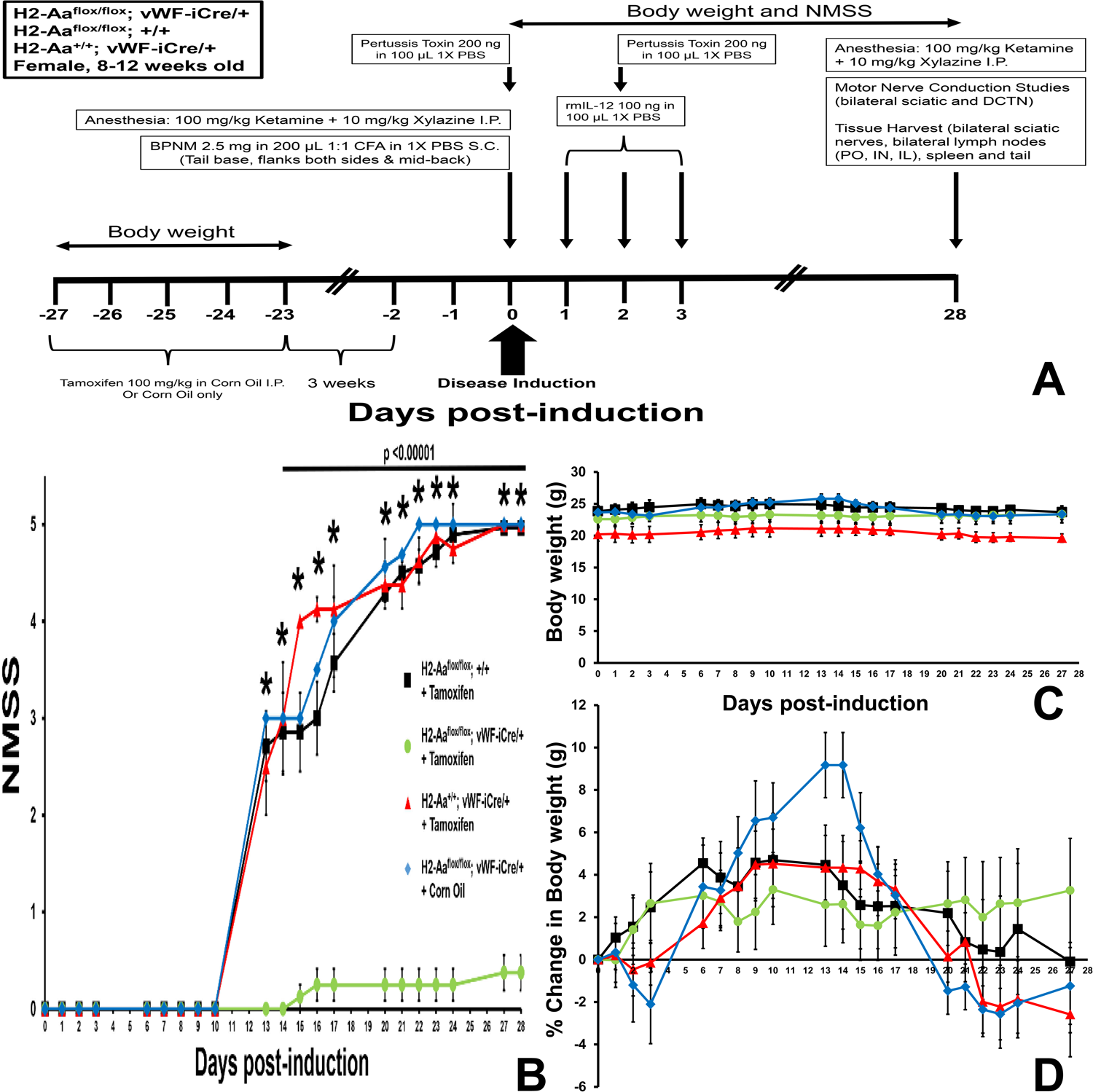
Sm-EAN experimental design and effect of sm-EAN on NMSS and weights in Tamoxifen-treated female SJL H2-Aa^flox/flox^; vWF-iCre/+, H2-Aa^flox/flox^; +/+ and H2-Aa^+/+^; vWF-iCre/+ mice and corn oil only-treated SJL H2-Aa^flox/flox^; vWF-iCre/+ mice. The experimental strategy used to induce and evaluate sm-EAN in experimental mice is shown in **A**. Microvascular endothelial cell H2-Aa knockout is associated with resistance to sm-EAN based on NMSS (**B**, p <0.001 from day 13, and p <0.0001 for the series from day 13 to 28 post-induction, Mann-Whitney U-test), with no significant differences in mean body weight (**C**) or % mean change in body weight (**D**) between experimental mice (p >0.05, ANOVA). DCTN: dorsal caudal tail nerve, IL = iliac, IN= inguinal, PO = popliteal, rmIL-12: recombinant mouse IL-12. Error bars indicate standard errors of the means. N= 8 for Tamoxifen-treated H2-Aa^flox/flox^; vWF-iCre/+, 7 for Tamoxifen-treated H2-Aa^flox/flox^; +/+ and 4 for both Tamoxifen-treated H2-Aa^+/+^; vWF-iCre/+ and corn oil only-treated SJL H2-Aa^flox/flox^; vWF-iCre/+ mice.

Sciatic motor nerve electrophysiology at expected sm-EAN peak severity showed significantly higher mean compound motor action potential (CMAP) amplitudes, faster conduction velocities, shorter waveform durations and shorter distal latencies in Tamoxifen-treated female SJL H2-Aa^flox/flox^; vWF-iCre/+ compared to mice with unaltered microvascular endothelial cell H2-Aa gene expression (**Figure 4**), consistent with NMSS scores. Similarly, DCT motor nerve electrophysiology at sm-EAN peak severity demonstrated similar findings to the sciatic nerve data, with the exception of the mean distal latencies (**Figure 5**). Since distal latency is dependent on terminal axon conduction and the neuromuscular junction conduction delay times following electrical stimulation, proximal tail histological evaluation to look for differences between Tamoxifen-treated sm-EAN-resistant female SJL H2-Aa^flox/flox^; vWF-iCre/+ and sm-EAN-affected SJL H2-Aa^flox/flox^; +/+ was performed. Representative motor nerve electrophysiology waveforms are shown in **Supplementary Figure 3**.

**Figure 4.**
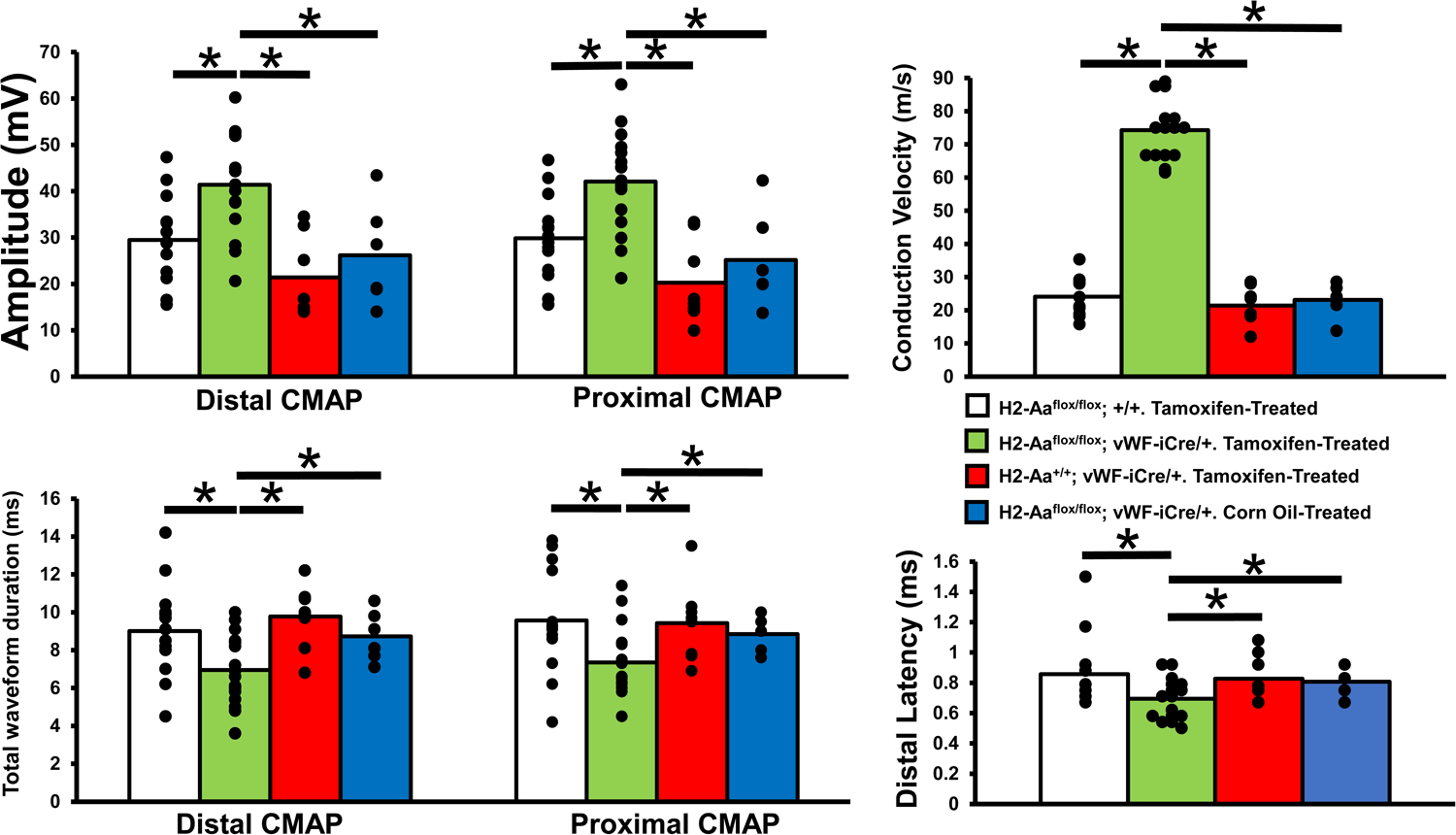
Sciatic nerve motor electrophysiology in Tamoxifen-treated female SJL H2-Aa^flox/flox^; vWF-iCre/+, H2-Aa^flox/flox^; +/+, and H2-Aa^+/+^; vWF-iCre/+ mice and corn oil only-treated SJL H2-Aa^flox/flox^; vWF-iCre/+ mice. Bar histograms show significantly higher mean distal and proximal CMAP amplitudes (indicative of axon numbers and integrity) and conduction velocity (indicative of myelination status of the fastest conducting axons) in microvascular endothelial cell H2-Aa knockout mice. Significantly reduced mean distal latency (indicative of distal axon and neuromuscular junction conduction delay time) and distal and proximal CMAP waveform durations (indicative of myelinated axon conduction synchrony) is seen in these mice compared to control mice with unaltered microvascular endothelial cell H2-Aa gene expression that fully develop electrophysiological features of a severe demyelinating neuritis with secondary axonal loss, consistent with sm-EAN. * indicate p < 0.05, ANOVA. Individual data points are shown for each sciatic nerve studied. N = 8 for Tamoxifen-treated H2-Aa^flox/flox^; vWF-iCre/+, 7 for Tamoxifen-treated H2-Aa^flox/flox^; +/+ and 4 for both Tamoxifen-treated H2-Aa^+/+^; vWF-iCre/+ and corn oil only-treated SJL H2-Aa^flox/flox^; vWF-iCre/+ mice.

**Figure 5.**
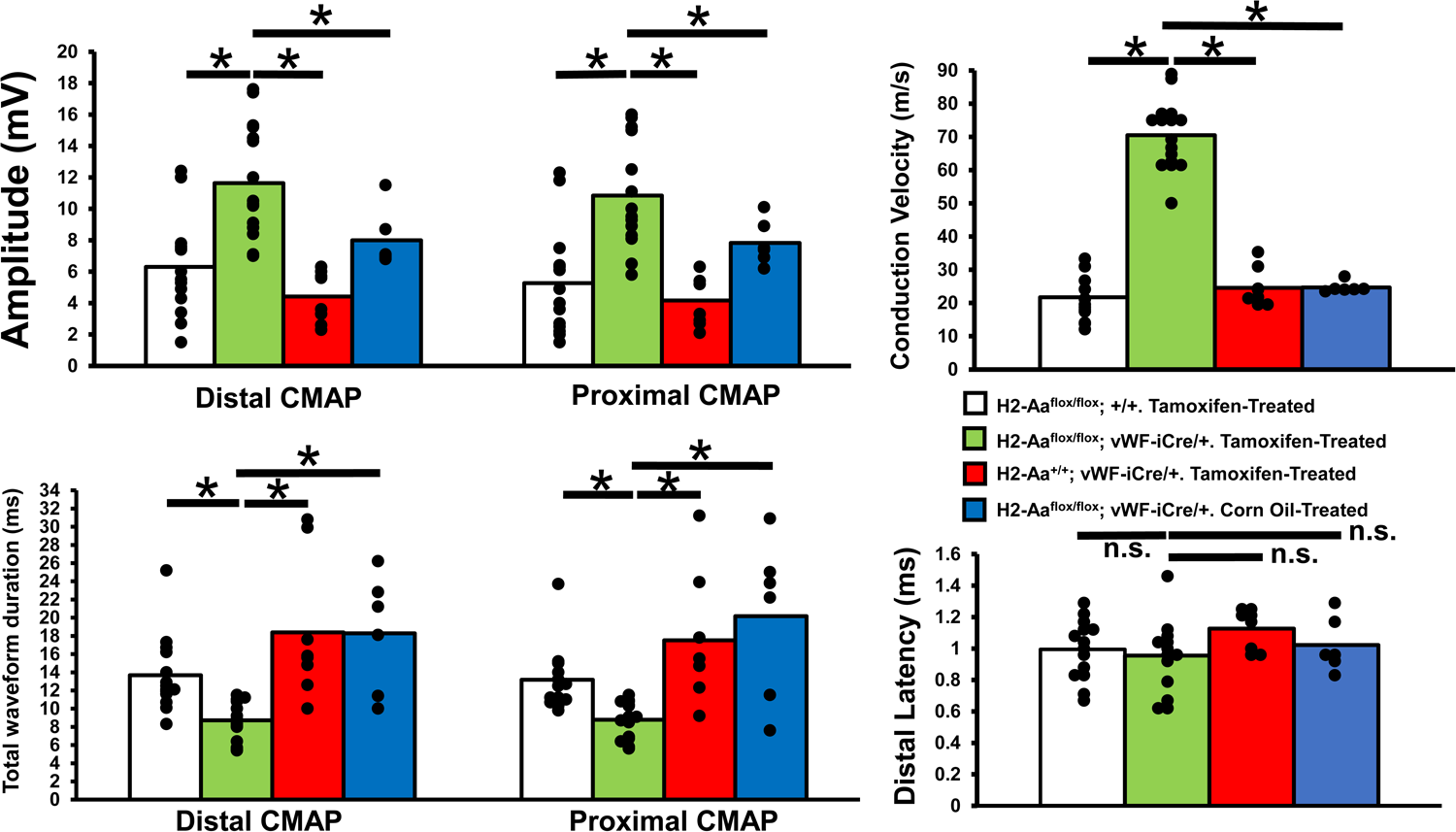
DCT nerve motor electrophysiology in Tamoxifen-treated female SJL H2-Aa^flox/flox^; vWF-iCre/+, H2-Aa^flox/flox^; +/+, and H2-Aa^+/+^; vWF-iCre/+ mice and corn oil only-treated SJL H2-Aa^flox/flox^; vWF-iCre/+ mice. Bar histograms show significantly higher mean distal and proximal CMAP amplitudes and conduction velocity in microvascular endothelial cell H2-Aa knockout mice. Significantly reduced mean distal and proximal waveform durations are seen in these mice compared to age-matched controls without Cre recombinase-mediated microvascular endothelial cell H2-Aa gene deletion that fully develop electrophysiological features of a severe demyelinating neuritis with secondary axonal loss, consistent with sm-EAN. No significant difference in mean distal latency is seen in experimental mice. * indicates p < 0.05, and n.s. = not significant (p > 0.05, ANOVA). Individual data points are shown for each DCT nerve studied. N = 8 for Tamoxifen-treated H2-Aa^flox/flox^; vWF-iCre/+, 7 for Tamoxifen-treated H2-Aa^flox/flox^; +/+ and 4 for both Tamoxifen-treated H2-Aa^+/+^; vWF-iCre/+ mice and corn oil only-treated SJL H2-Aa^flox/flox^; vWF-iCre/+ mice.

Semi-thin plastic-embedded sections at expected sm-EAN peak severity showed normal sciatic nerves in Tamoxifen-treated female SJL H2-Aa^flox/flox^; vWF-iCre/+ compared to inflammatory demyelination with axonal loss observed in female SJL H2-Aa^flox/flox^; +/+ mice (**Figure 6**). Interestingly, leukocyte infiltration was observed in extraneural connective tissue in sm-EAN-resistant H2-Aa^flox/flox^; vWF-iCre/+ mice, suggesting that effector leukocyte transmigration directional cues for peripheral nerve infiltration was altered following abrogated microvascular endothelial cell MHC class II expression. Indirect fluorescent immunohistochemistry at expected sm-EAN peak severity showed absent or rare sciatic nerve H2-A+ CD45+ leukocytes in Tamoxifen-treated female SJL H2-Aa^flox/flox^; vWF-iCre/+ mice compared to intense infiltration observed in SJL H2-Aa^flox/flox^; +/+ mice, indicative of inhibited inflammation (**Figure 7 and Supplementary Figure 4**). Interestingly, perineurial H2-A expression was observed in Tamoxifen-treated female SJL H2-Aa^flox/flox^; vWF-iCre/+ mice, and increased diffuse H2-A expression was observed in female SJL H2-Aa^flox/flox^; +/+ mice with sm-EAN, consistent with AIDP-GBS. These findings were also observed in DCT nerves (**Figure 7 and Supplementary Figure 4**); however, H2-A expression was similarly observed in coccygeal bone marrow leukocyte clusters between Tamoxifen-treated female SJL H2-Aa^flox/flox^; vWF-iCre/+ and SJL H2-Aa^flox/flox^; +/+ mice. Cutaneous inflammation consisting of H2-A+ and H2-A-CD45+ leukocytes occurred in Tamoxifen-treated H2-Aa^flox/flox^; vWF-iCre/+ mice with tail weakness <u>without</u> DCT nerve inflammation (**Figure 7 and Supplementary Figure 4**), providing an explanation for the observed NMSS and prolonged mean distal latencies. Sm-EAN peak severity indirect fluorescent immunohistochemistry showed H2-A expression by spleen and inguinal lymph node CD45+ leukocyte subsets in Tamoxifen-treated female SJL H2-Aa^flox/flox^; vWF-iCre/+ mice (**Figure 8**). This provided evidence that microvascular endothelial cell-specific MHC class II deletion did not inadvertently alter professional APC organization in secondary lymphoid organs. Splenomegaly, with disorganized H2-A+ CD45+ leukocyte architecture was observed in sm-EAN-affected Tamoxifen-treated H2-Aa^flox/flox^; +/+ mice, as previously described in mice with active autoimmune disease.^55,56^

**Figure 6.**
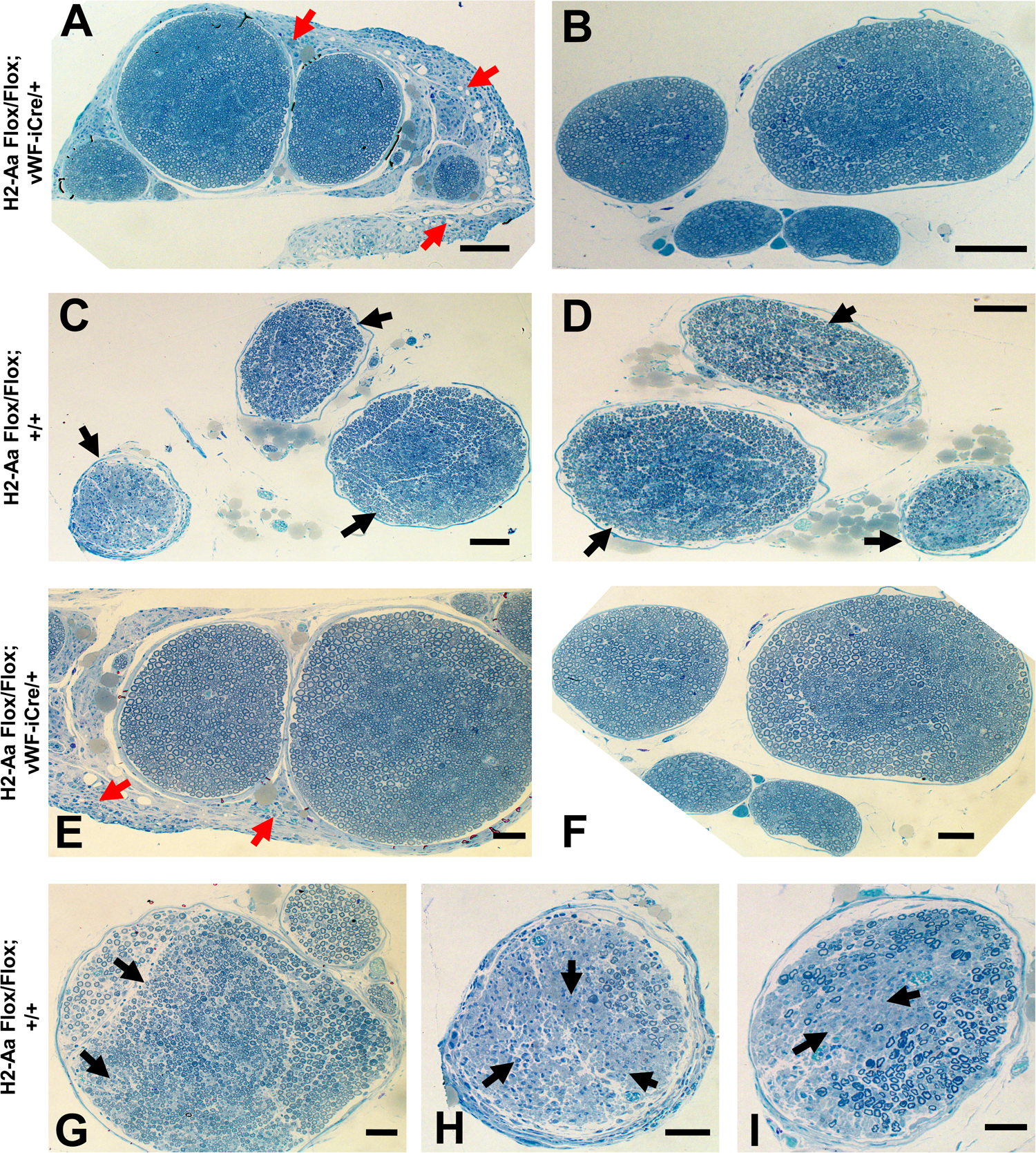
Comparative sciatic nerve morphology in Tamoxifen-treated female SJL H2-Aa^flox/flox^; vWF-iCre/+ and H2-Aa^flox/flox^; +/+ mice. Representative digital photomicrographs of Toluidine blue-stained semi-thin plastic-embedded axial sections show normal sciatic nerve morphology in microvascular endothelial cell H2-Aa knockout mice at low (**A, B**) and higher (**E, F**) magnification, with leukocyte infiltrates seen in the epineurial and extraneural connective tissue (**A and E**, red arrows). This is in contrast to inflammatory cell infiltration, demyelination and axonal loss (black arrows) seen in sm-EAN-affected mice without microvascular endothelial cell H2-Aa gene deletion at low (**C-D**) and higher (**G-I**) magnification. Scale bars: A-D = 200 μm, E-I = 100 μm.

**Figure 7.**
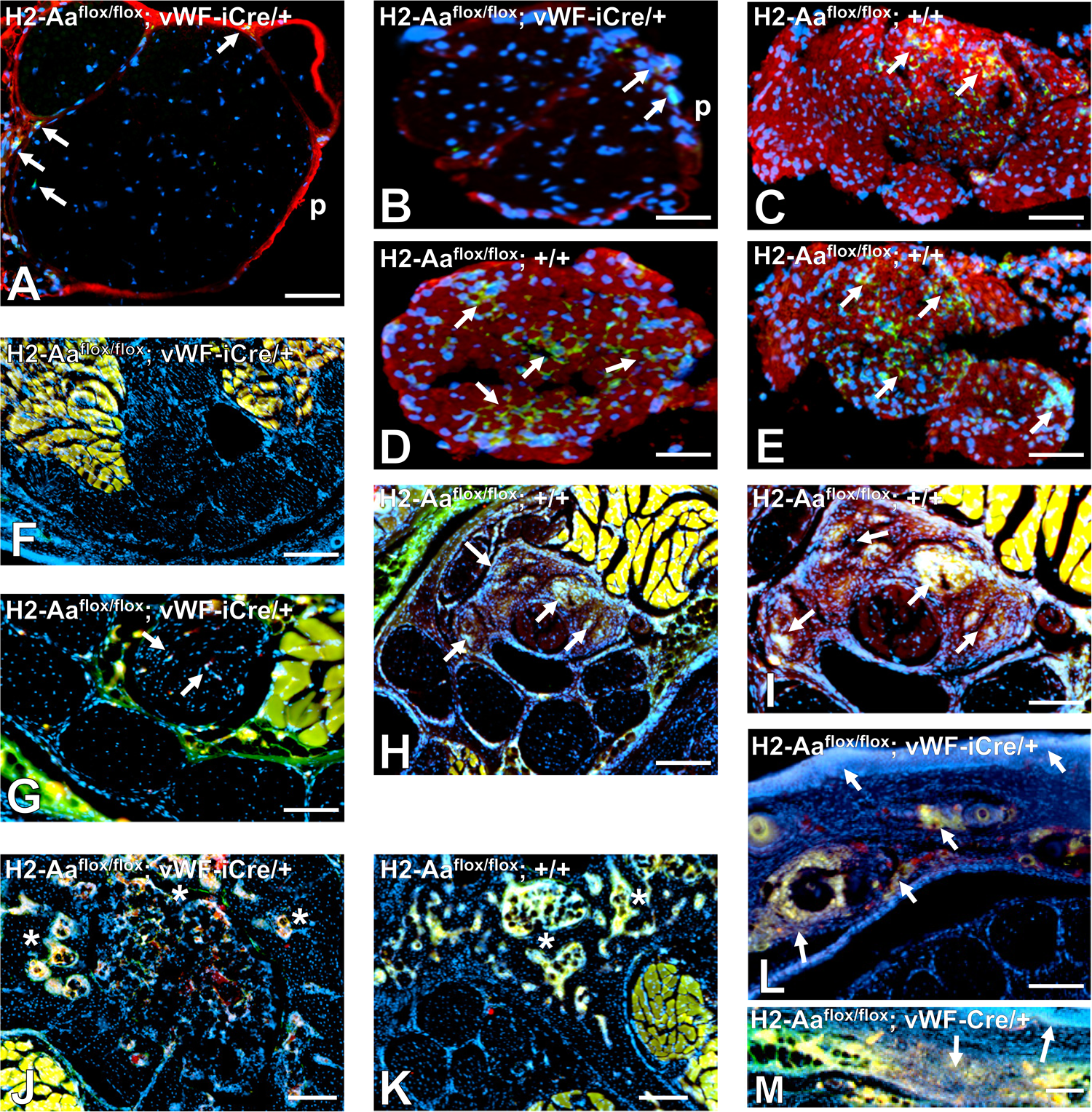
Comparative sciatic nerve and tail indirect fluorescent immunohistochemistry in Tamoxifen-treated female SJL H2-Aa^flox/flox^; vWF-iCre/+ and H2-Aa^flox/flox^; +/+ mice. Representative digital photomicrographs show perineurial (p) and rare perineurial and endoneurial H2-A+ and H2-A-CD45+ leukocytes in the sciatic nerves of microvascular endothelial cell H2-Aa knockout mice (**A-B**, white arrows). This is in contrast to diffusely increased H2-A expression and multifocal endoneurial H2-A+ CD45+ leukocyte infiltration observed in sm-EAN affected mice without H2-Aa gene deletion (**C-E**, white arrows). DCT nerve endoneurial H2-A+ CD45+ leukocyte infiltration is minimal in a microvascular endothelial cell H2-Aa knockout mouse at low (**F**) and higher (**G,** white arrows) magnification, in contrast to diffusely increased H2-A and multifocal endoneurial H2-A+ CD45+ leukocyte infiltration (white arrows) in an sm-EAN affected mouse without H2-Aa gene deletion at low (**H**) and higher (**I**) magnification. Clusters of coccygeal bone marrow H2-A+ CD45+ leukocytes (asterisk) are seen in both microvascular endothelial cell H2-Aa knockout mice (**J**) and mice without H2-Aa gene deletion (**K**). Epidermal and dermal inflammation consisting mainly of H2-A+ CD45+ leukocytes is seen in a microvascular endothelial cell H2-Aa knockout mouse with tail weakness without DCT nerve inflammation (**L-M**, white arrows). Local extraneural tissue inflammation with edema provides an explanation for similar mean DCT nerve distal latencies in microvascular endothelial cell H2-Aa knockout mice and sm-EAN-affected mice without H2-Aa gene deletion. Non-specific red and green fluorescence is seen in all tail muscle fibers. Scale bars: A-E = 50 μm, F and H = 100 μm, G and I = 50 μm, J-M = 100 μm.

**Figure 8.**
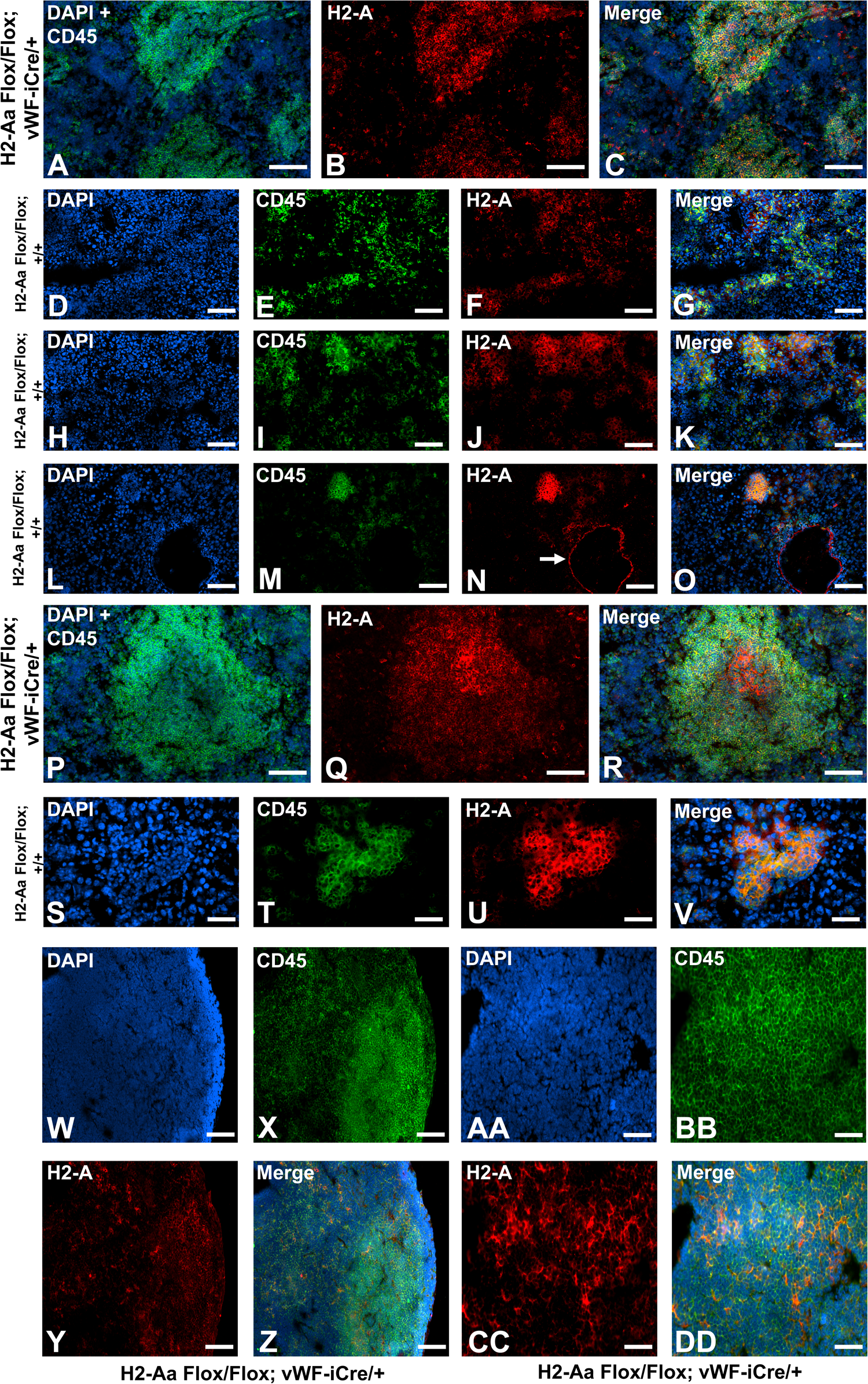
Comparative spleen and inguinal lymph node indirect fluorescent immunohistochemistry in Tamoxifen-treated female SJL H2-Aa^flox/flox^; vWF-iCre/+ and H2-Aa^flox/flox^; +/+ mice. Representative digital photomicrographs from the spleen of a microvascular endothelial cell H2-Aa knockout mouse show organized large foci of H2-A+ and H2-A-CD45+ leukocytes at low (**A-C**) and higher (**P-R**) magnification. Disorganized diffuse and small foci of H2-A+ CD45+ leukocytes are seen in an sm-EAN-affected mouse without H2-Aa gene deletion with massive splenomegaly (**D-K**). Disorganized, but larger foci of H2-A+ CD45+ leukocytes are seen in an sm-EAN-affected mouse without H2-Aa gene deletion with splenomegaly at low (**L-O**) and higher (**S-V**) magnification. H2-A+ spleen sinusoidal endothelial cells in an sm-EAN affected mouse without H2-Aa gene deletion with splenomegaly (white arrow) is shown in **N**. Diffuse H2-Aa+ and H2-Aa-CD45+ leukocytes are seen in the inguinal lymph node of a microvascular endothelial cell H2-Aa knockout mouse resistant to sm-EAN (**W-DD**), as seen in sm-EAN affected mice without H2-Aa gene deletion. Scale bars: A-O = 100 μm, P-V = 50 μm, W-Z = 100 μm, AA-DD = 50 μm.

## Discussion

Our initial studies show that microvascular endothelial cell-specific H2-Aa knockout prevented acute autoimmune demyelinating neuritis in female adult SJL mice, implying that BNB-forming endothelial cell MHC Class II expression is necessary for antigen-specific peripheral nerve-restricted autoimmunity. Comparative quantitative histopathological analyses were not performed due to the observed “all-or-none” differences in inflammatory demyelination with axonal loss in sm-EAN-affected mice with unaltered microvascular endothelial cell H2-Aa gene expression compared to Tamoxifen-treated SJL H2-Aa^flox/flox^; vWF-iCre/+ mice with microvascular endothelial cell-specific H2-Aa gene deletion. This supports human *in vitro* data that demonstrated vascular endothelial cell capacity to promote antigen presentation, effector memory T-cell proliferation, and antigen-specific CD4+ T-cell migration.^1,22,24–26^ Importantly, leukocyte H2-A expression was not unexpectedly altered in Tamoxifen-treated H2-Aa^flox/flox^; vWF-iCre/+ SJL mouse bone marrow or secondary lymphoid organs to suggest altered professional APC function required to initiate adaptive immune responses to exogenous antigens. Despite the limitations of human autoimmune disease murine models, observations from this study provide pathogenic insights into human AIDP due to similarities with sm-EAN.

Sm-EAN reliably occurs in 100% of female SJL mice and requires subcutaneous BPNM-CFA <u>and</u> Pertussis toxin, with more severe disease observed with recombinant mouse IL-12 as a co-adjuvant.^46^ Histological characterization of inflammatory demyelination with axonal loss in sm-EAN has been previously published.^13,47,49^ BPNM-CFA alone is insufficient to induce sm-EAN and Pertussis toxin has been shown to potently activate MHC Class II expression in lymphoid and nonlymphoid target organs and induce proliferation of antigen-specific CD4+ T-helper (Th) 1 and Th2 cells in addition to histamine- and cytokine-mediated increased microvascular permeability.^57,58^ Recombinant mouse IL-12 promotes differentiation of naïve CD4+ T-cells to CD4+ Th1 cells. MHC class II upregulation occurred in BNB-forming endoneurial endothelial cells at sm-EAN onset and was associated with leukocyte transmigration *in situ*. Diffuse MHC class II expression occurred in sm-EAN at peak severity, as seen in AIDP-GBS. Sural nerve biopsy from AIDP-GBS patients at disease onset is not ethically feasible to validate sm-EAN data. However, MHC class II was constitutively expressed by histologically normal adult endoneurial endothelial cells to suggest an important role in normal immune surveillance.

SJL mouse adaptive immune responses require H2-Aa, as these mice are H2-Ea null and redundancy exists for β-chain molecules.^38^ H2-Aa (with human homolog HLA-DQA1) is highly conserved between species, suggesting an essential role in adaptive immunity. Our construct selectively excises exons 2, 3, 4 and the 5’ portion of exon 5 in Cre recombinase-expressing cells, generating a nonfunctional H2-Aa protein. Polymorphisms (e.g. at exon 2) could explain differential susceptibility to sm-EAN between SJL and C57BL/6 mice. Although conclusive HLA class II susceptibility risk genes have not been identified in GBS,^59–62^ HLA-DQA1 is a plausible target for deep sequencing in worldwide AIDP-GBS patient cohorts based on sm-EAN data.

Further work is required to confirm these initial results, determine whether delayed inflammation occurs with antigen persistence >28 days and confirm normal MHC Class II antigen presentation cell function in sm-EAN-resistant Tamoxifen-treated SJL H2-Aa^flox/flox^; vWF-iCre/+ mice. The molecular determinants and signaling mechanisms required for antigen-specific effector memory CD4+ T-cell engagement with endoneurial endothelial cells via MHC class II as a prerequisite for pathogenic leukocyte transmigration and acute demyelinating neuritis in sm-EAN, with potential relevance to immune-mediated polyneuropathies, needs to be elucidated.

The conditional H2-Aa knockout mouse strains provide essential tools to study antigen presentation by different cells during tissue- or organ-specific adaptive immune responses in a time-dependent manner. Tamoxifen-induced Cre recombination effectively and maximally excises the floxed H2-Aa gene three weeks after 5 consecutive daily injections,^45^ facilitating timed gene deletion to disease model phase. Tissue or organ grafts from mice with microvascular endothelial cell-specific MHC class II knockout could be used to study allograft and xenograft rejection *in vivo*. Studies using living mice with intact immune systems are advantageous over human *in vitro* and *ex vivo* studies, provided such studies are guided by patient observational data. Improved understanding of molecular mimicry, tissue-specific autoimmunity and transplant rejection obtained from conditional MHC class II knockout mouse studies should advance knowledge and guide targeted immune drug development strategies.

## Acknowledgments

The H2-Aa SJL mouse strain was developed from embryonic stem (ES) cell clone H2-Aa G03 (EPD0585_5_G03), obtained from the University of California Davis Knockout Mouse Project (KOMP) Repository (www.komp.org) and generated by the Wellcome Trust Sanger Institute. Targeting vectors used were generated by the Wellcome Trust Sanger Institute and the Children’s Hospital Oakland Research Institute as part of KOMP (Grant # 3U01HG004080).

We acknowledge Elizabeth Hughes for culturing the H2-Aa^tm1a(KOMP)Wtsi^ Targeted ES cell clone and Galina Gavrilina for blastocyst microinjection to produce the H2-Aa ES cell-mouse chimeras, and the Transgenic Animal Model Core of the University of Michigan’s Biomedical Research Core Facilities for their excellent technical expertise. Special thanks to past members of the Neuromuscular Immunopathology Research Laboratory, particularly Dr. Chaoling Dong, for their essential technical assistance required to develop, characterize and maintain the conditional H2-Aa C57BL/6 and SJL mouse colonies at the University of Alabama at Birmingham (UAB). We acknowledge the University of Missouri MMRRC for validating our transgenic mouse strains and performing transgene copy number analysis on our inducible von Willebrand factor Cre recombinase SJL mouse strain. Special thanks to Beth Weeks (EMLabs Inc.) for technical assistance with sciatic nerve plastic embedding, semi-thin sectioning and staining. Thanks for Dr. Jordan Pober for insightful discussions on human microvascular endothelial cell antigen presentation *in vitro*.

## Ethical Publication Statement

We confirm that we have read the Journal’s position on issues involved in ethical publication and affirm that this report is consistent with those guidelines

## Data Availability Statement

Data that support the study’s findings are available to members of the scientific community on request from the corresponding author, E.E.U.

## Funding Statement

This work was supported by institutional funds from the University of Alabama at Birmingham to E.E.U.

## Conflict of Interest Disclosure

E.E.U. has received research support from the National Institute of Neurological Disorders and Stroke/ National Institutes of Health, CSL Behring, Sanofi-Genzyme and Argenx BBVA, and royalties from a non-exclusive commercial license for a simian virus 40 large T-cell immortalized human endoneurial endothelial cell line (held by Baylor College of Medicine Ventures) and books published by Springer Science + Business Media. E.E.U. has also served as a paid consultant for UCB, Latigo Biotherapeutics and Seismic Therapeutics, as well as a paid grant reviewer for the Peer Reviewed Medical Research Program and Paul G. Allen Frontiers Group during the period of the research activity and generation of the current report. The C57BL/6 H2-Aa conditional knockout (C57BL/6-*H2-Aa^tm1c(KOMP)Wist^*/UbeeMmmh; RRID:MMRRC_046053-MU) SJL H2-Aa conditional knockout (SJL.B6-*H2-Aa^tm1c(KOMP)Wtsi^*/UbeeMmmh; RRID:MMRRC_071292-MU) and Tamoxifen-inducible von Willebrand factor (SJL.Cg-Tg(VWF-Cre/ERT2)C1014/UbeeMmmh; RRID:MMRRC_071293-MU) transgenic mouse strains are housed and commercially available from the University of Missouri Mutant Mouse Resource & Research Center (MMRRC) for worldwide distribution to academic, non-profit and for-profit organizations and companies based on Materials Transfer Agreements and commercial licensing contracts with the University of Alabama at Birmingham Harbert Institute for Innovation and Entrepreneurship (HIEE). The remaining authors have nothing to disclose

## Ethics Approval Statements

The use of archived de-identified patient sural nerve biopsies from the UAB Hospital Shin J. Oh Muscle and Nerve Histopathology Laboratory for biomedical research was approved by the Institutional Review Board (Protocol #140321012) and subsequently determined exempt from annual review, based on the 2018 Revised Common Rule regulations (Category 4), as of 11/22/2019.

All experimental mouse studies were conducted in accordance with all provisions of the Animal Welfare Act, the U.S. Government Principles Regarding the Care and Use of Animals, the Guide (Guide for the Care and Use of Laboratory Animals), National Research Council, Institute of Laboratory Animal Resources, the Division of Public Health Service Policy on Humane Care and Use of Laboratory Animals and the additional local policies of the Center of Comparative Medicine, UAB under approved Institutional Animal Care and Use Committee (Protocol # 141009960).

## Abbreviations

AIDP: acute inflammatory demyelinating polyradiculoneuropathy
ANOVA: analysis of variance
BNB: blood-nerve barrier
bp: base pairs
BPNM: bovine peripheral nerve myelin
C57BL/6: C57 Black 6
CCD: charged-coupled device
CD: Cluster of differentiation
CFA: Complete Freund Adjuvant
CMAP: compound motor action potential
DAPI: 4′,6-diamidino-2-phenylindole
DCT: dorsal caudal tail
DNA: deoxyribonucleic acid
dNTP: deoxyribonucleotide triphosphate
D-PBS: Dulbecco’s phosphate buffered saline
EDTA: Ethylenediaminetetraacetic acid
ES: embryonic stem
flpo: flippase o
FITC: fluorescein isothiocyanate
FRT: flippase recombinase target
GBS: Guillain-Barré syndrome
GTA: glutaraldehyde
H2-Aa: mouse Major Histocompatibility Complex class II antigen A-alpha chain
HLA: human leukocyte antigen
IL: interleukin
loxP: “locus of crossing over in P1”
MHC: Major Histocompatibility Complex
MMRRC: Mutant Mouse Resource & Research Center
NGS: normal goat serum
NMSS: Neuromuscular Severity Score Scale
OCT: Optimum Cutting Temperature^®^
PBS: phosphate buffered saline
PFA: paraformaldehyde
PCR: polymerase chain reaction
SJL: Swiss Jim Lambert
Sm-EAN: severe murine experimental autoimmune neuritis
SNP: single nucleotide polymorphism
RRID: Research Resource Identifier
RT: room temperature
TAE: Tris base, Acetic acid, Ethylenediaminetetraacetic acid
tdT: tdTomato
Th: T-helper
TL: Tomato lectin
UAB: University of Alabama at Birmingham
UEA-1: *Ulex Europaeus Agglutinin-1*
vWF: von Willebrand factor
vWF-iCre: inducible von Willebrand factor Cre recombinase transgene

## Notes

### Competing Interest Statement

The authors have declared no competing interest.

### Summary of Updates

The manuscript, figures and Supplemental files have been significantly changed in response to scientific peer-review.

## References

1. Pober JS, Merola J, Liu R, Manes TD. Antigen Presentation by Vascular Cells. Front Immunol 2017;8:1907.

2. Rosenblum MD, Remedios KA, Abbas AK. Mechanisms of human autoimmunity. The Journal of clinical investigation 2015;125(6):2228–2233.

3. Rojas M, Restrepo-Jimenez P, Monsalve DM, Pacheco Y, Acosta-Ampudia Y, Ramirez-Santana C, Leung PSC, Ansari AA, Gershwin ME, Anaya JM. Molecular mimicry and autoimmunity. J Autoimmun 2018;95:100–123.

4. Gray JI, Westerhof LM, MacLeod MKL. The roles of resident, central and effector memory CD4 T-cells in protective immunity following infection or vaccination. Immunology 2018;154(4):574–581.

5. Woodland DL, Kohlmeier JE. Migration, maintenance and recall of memory T cells in peripheral tissues. Nat Rev Immunol 2009;9(3):153–161.

6. van den Berg B, Walgaard C, Drenthen J, Fokke C, Jacobs BC, van Doorn PA. Guillain-Barre syndrome: pathogenesis, diagnosis, treatment and prognosis. Nature reviews Neurology 2014;10(8):469–482.

7. Willison HJ, Jacobs BC, van Doorn PA. Guillain-Barre syndrome. Lancet 2016;388(10045):717–727.

8. Hughes RAC, Cornblath DR, Willison HJ. Guillain-Barre syndrome in the 100 years since its description by Guillain, Barre and Strohl. Brain : a journal of neurology 2016;139(11):3041–3047.

9. Hughes R, Atkinson P, Coates P, Hall S, Leibowitz S. Sural nerve biopsies in Guillain-Barre syndrome: axonal degeneration and macrophage-associated demyelination and absence of cytomegalovirus genome. Muscle & nerve 1992;15(5):568–575.

10. Nyland H, Matre R, Mork S. Immunological characterization of sural nerve biopsies from patients with Guillain-Barre syndrome. Annals of neurology 1981;9 Suppl:80-86.

11. Putzu GA, Figarella-Branger D, Bouvier-Labit C, Liprandi A, Bianco N, Pellissier JF. Immunohistochemical localization of cytokines, C5b-9 and ICAM-1 in peripheral nerve of Guillain-Barre syndrome. Journal of the neurological sciences 2000;174(1):16–21.

12. Schmidt B, Toyka KV, Kiefer R, Full J, Hartung HP, Pollard J. Inflammatory infiltrates in sural nerve biopsies in Guillain-Barre syndrome and chronic inflammatory demyelinating neuropathy. Muscle & nerve 1996;19(4):474–487.

13. Dong C, Palladino SP, Helton ES, Ubogu EE. The pathogenic relevance of alphaM-integrin in Guillain-Barre syndrome. Acta neuropathologica 2016;132(5):739–752.

14. Ubogu EE. Inflammatory neuropathies: pathology, molecular markers and targets for specific therapeutic intervention. Acta neuropathologica 2015;130(4):445–468.

15. Ley K, Laudanna C, Cybulsky MI, Nourshargh S. Getting to the site of inflammation: the leukocyte adhesion cascade updated. Nat Rev Immunol 2007;7(9):678–689.

16. Man S, Ubogu EE, Ransohoff RM. Inflammatory cell migration into the central nervous system: a few new twists on an old tale. Brain pathology 2007;17(2):243–250.

17. Muller WA. Leukocyte-endothelial-cell interactions in leukocyte transmigration and the inflammatory response. Trends Immunol 2003;24(6):327–334.

18. Muller WA. Mechanisms of leukocyte transendothelial migration. Annu Rev Pathol 2011;6:323–344.

19. Muller WA. How endothelial cells regulate transmigration of leukocytes in the inflammatory response. Am J Pathol 2014;184(4):886–896.

20. Sathaliyawala T, Kubota M, Yudanin N, Turner D, Camp P, Thome JJ, Bickham KL, Lerner H, Goldstein M, Sykes M, Kato T, Farber DL. Distribution and compartmentalization of human circulating and tissue-resident memory T cell subsets. Immunity 2013;38(1):187–197.

21. Manes TD, Pober JS. Identification of endothelial cell junctional proteins and lymphocyte receptors involved in transendothelial migration of human effector memory CD4+ T cells. J Immunol 2011;186(3):1763–1768.

22. Manes TD, Pober JS. Antigen presentation by human microvascular endothelial cells triggers ICAM-1-dependent transendothelial protrusion by, and fractalkine-dependent transendothelial migration of, effector memory CD4+ T cells. J Immunol 2008;180(12):8386–8392.

23. Rothermel AL, Wang Y, Schechner J, Mook-Kanamori B, Aird WC, Pober JS, Tellides G, Johnson DR. Endothelial cells present antigens in vivo. BMC Immunol 2004;5:5.

24. Wheway J, Obeid S, Couraud PO, Combes V, Grau GE. The brain microvascular endothelium supports T cell proliferation and has potential for alloantigen presentation. PloS one 2013;8(1):e52586.

25. Lopes Pinheiro MA, Kamermans A, Garcia-Vallejo JJ, van Het Hof B, Wierts L, O’Toole T, Boeve D, Verstege M, van der Pol SM, van Kooyk Y, de Vries HE, Unger WW. Internalization and presentation of myelin antigens by the brain endothelium guides antigen-specific T cell migration. Elife 2016;5.

26. Manes TD, Wang V, Pober JS. Costimulators expressed on human endothelial cells modulate antigen-dependent recruitment of circulating T lymphocytes. Front Immunol 2022;13:1016361.

27. Abrahimi P, Qin L, Chang WG, Bothwell AL, Tellides G, Saltzman WM, Pober JS. Blocking MHC class II on human endothelium mitigates acute rejection. JCI Insight 2016;1(1).

28. Van Rhijn I, Van den Berg LH, Bosboom WM, Otten HG, Logtenberg T. Expression of accessory molecules for T-cell activation in peripheral nerve of patients with CIDP and vasculitic neuropathy. Brain : a journal of neurology 2000;123 (Pt 10):2020–2029.

29. Matsumuro K, Izumo S, Umehara F, Osame M. Chronic inflammatory demyelinating polyneuropathy: histological and immunopathological studies on biopsied sural nerves. Journal of the neurological sciences 1994;127(2):170–178.

30. Mitchell GW, Williams GS, Bosch EP, Hart MN. Class II antigen expression in peripheral neuropathies. Journal of the neurological sciences 1991;102(2):170–176.

31. Meyer Zu Horste G, Heidenreich H, Lehmann HC, Ferrone S, Hartung HP, Wiendl H, Kieseier BC. Expression of antigen processing and presenting molecules by Schwann cells in inflammatory neuropathies. Glia 2010;58(1):80–92.

32. Pollard JD, Baverstock J, McLeod JG. Class II antigen expression and inflammatory cells in the Guillain-Barre syndrome. Annals of neurology 1987;21(4):337–341.

33. Pollard JD, McCombe PA, Baverstock J, Gatenby PA, McLeod JG. Class II antigen expression and T lymphocyte subsets in chronic inflammatory demyelinating polyneuropathy. Journal of neuroimmunology 1986;13(2):123–134.

34. Meyer zu Horste G, Heidenreich H, Mausberg AK, Lehmann HC, ten Asbroek AL, Saavedra JT, Baas F, Hartung HP, Wiendl H, Kieseier BC. Mouse Schwann cells activate MHC class I and II restricted T-cell responses, but require external peptide processing for MHC class II presentation. Neurobiol Dis 2010;37(2):483–490.

35. Hartlehnert M, Derksen A, Hagenacker T, Kindermann D, Schafers M, Pawlak M, Kieseier BC, Meyer Zu Horste G. Schwann cells promote post-traumatic nerve inflammation and neuropathic pain through MHC class II. Sci Rep 2017;7(1):12518.

36. Palladino SP, Helton ES, Jain P, Dong C, Crowley MR, Crossman DK, Ubogu EE. The Human Blood-Nerve Barrier Transcriptome. Sci Rep 2017;7(1):17477.

37. Gerber D, Pereira JA, Gerber J, Tan G, Dimitrieva S, Yanguez E, Suter U. Transcriptional profiling of mouse peripheral nerves to the single-cell level to build a sciatic nerve ATlas (SNAT). Elife 2021;10.

38. Monzon-Casanova E, Rudolf R, Starick L, Muller I, Sollner C, Muller N, Westphal N, Miyoshi-Akiyama T, Uchiyama T, Berberich I, Walter L, Herrmann T. The Forgotten: Identification and Functional Characterization of MHC Class II Molecules H2-Eb2 and RT1-Db2. J Immunol 2016;196(3):988–999.

39. Kranz A, Fu J, Duerschke K, Weidlich S, Naumann R, Stewart AF, Anastassiadis K. An improved Flp deleter mouse in C57Bl/6 based on Flpo recombinase. Genesis 2010;48(8):512–520.

40. Jahroudi N, Lynch DC. Endothelial-cell-specific regulation of von Willebrand factor gene expression. Mol Cell Biol 1994;14(2):999–1008.

41. Aird WC, Jahroudi N, Weiler-Guettler H, Rayburn HB, Rosenberg RD. Human von Willebrand factor gene sequences target expression to a subpopulation of endothelial cells in transgenic mice. Proc Natl Acad Sci U S A 1995;92(10):4567–4571.

42. Muzumdar MD, Tasic B, Miyamichi K, Li L, Luo L. A global double-fluorescent Cre reporter mouse. Genesis 2007;45(9):593–605.

43. Madisen L, Zwingman TA, Sunkin SM, Oh SW, Zariwala HA, Gu H, Ng LL, Palmiter RD, Hawrylycz MJ, Jones AR, Lein ES, Zeng H. A robust and high-throughput Cre reporting and characterization system for the whole mouse brain. Nat Neurosci 2010;13(1):133–140.

44. Hasegawa Y, Daitoku Y, Sekiguchi K, Tanimoto Y, Mizuno-Iijima S, Mizuno S, Kajiwara N, Ema M, Miwa Y, Mekada K, Yoshiki A, Takahashi S, Sugiyama F, Yagami K. Novel ROSA26 Cre-reporter knock-in C57BL/6N mice exhibiting green emission before and red emission after Cre-mediated recombination. Exp Anim 2013;62(4):295–304.

45. Jahn HM, Kasakow CV, Helfer A, Michely J, Verkhratsky A, Maurer HH, Scheller A, Kirchhoff F. Refined protocols of tamoxifen injection for inducible DNA recombination in mouse astroglia. Sci Rep 2018;8(1):5913.

46. Calida DM, Kremlev SG, Fujioka T, Hilliard B, Ventura E, Constantinescu CS, Lavi E, Rostami A. Experimental allergic neuritis in the SJL/J mouse: induction of severe and reproducible disease with bovine peripheral nerve myelin and pertussis toxin with or without interleukin-12. Journal of neuroimmunology 2000;107(1):1–7.

47. Xia RH, Yosef N, Ubogu EE. Clinical, electrophysiological and pathologic correlations in a severe murine experimental autoimmune neuritis model of Guillain-Barre syndrome. Journal of neuroimmunology 2010;219(1-2):54–63.

48. Xia RH, Yosef N, Ubogu EE. Selective expression and cellular localization of pro-inflammatory chemokine ligand/receptor pairs in the sciatic nerves of a severe murine experimental autoimmune neuritis model of Guillain-Barre syndrome. Neuropathology and applied neurobiology 2010;36(5):388–398.

49. Yuan F, Yosef N, Lakshmana Reddy C, Huang A, Chiang SC, Tithi HR, Ubogu EE. CCR2 gene deletion and pharmacologic blockade ameliorate a severe murine experimental autoimmune neuritis model of Guillain-Barre syndrome. PloS one 2014;9(3):e90463.

50. Xia RH, Yosef N, Ubogu EE. Dorsal caudal tail and sciatic motor nerve conduction studies in adult mice: technical aspects and normative data. Muscle & nerve 2010;41(6):850–856.

51. Dong C, Helton ES, Zhou P, Ouyang X, d’Anglemont de Tassigny X, Pascual A, Lopez-Barneo J, Ubogu EE. Glial-derived neurotrophic factor is essential for blood-nerve barrier functional recovery in an experimental murine model of traumatic peripheral neuropathy. Tissue Barriers 2018;6(2):1–22.

52. Yang D, Li S, Wu J. A Simple and Quick Method for Decalcification Using Mouse Tail as a Model for Preparation of Lymphedema Study. Appl Immunohistochem Mol Morphol 2021;29(7):551–556.

53. Brown D, Lydon J, McLaughlin M, Stuart-Tilley A, Tyszkowski R, Alper S. Antigen retrieval in cryostat tissue sections and cultured cells by treatment with sodium dodecyl sulfate (SDS). Histochem Cell Biol 1996;105(4):261–267.

54. Ouyang X, Dong C, Ubogu EE. In situ molecular characterization of endoneurial microvessels that form the blood-nerve barrier in normal human adult peripheral nerves. Journal of the peripheral nervous system : JPNS 2019;24(2):195–206.

55. Garchow BG, Bartulos Encinas O, Leung YT, Tsao PY, Eisenberg RA, Caricchio R, Obad S, Petri A, Kauppinen S, Kiriakidou M. Silencing of microRNA-21 in vivo ameliorates autoimmune splenomegaly in lupus mice. EMBO Mol Med 2011;3(10):605–615.

56. Goltsev Y, Samusik N, Kennedy-Darling J, Bhate S, Hale M, Vazquez G, Black S, Nolan GP. Deep Profiling of Mouse Splenic Architecture with CODEX Multiplexed Imaging. Cell 2018;174(4):968–981 e915.

57. Hofstetter HH, Shive CL, Forsthuber TG. Pertussis toxin modulates the immune response to neuroantigens injected in incomplete Freund’s adjuvant: induction of Th1 cells and experimental autoimmune encephalomyelitis in the presence of high frequencies of Th2 cells. J Immunol 2002;169(1):117–125.

58. Kugler S, Bocker K, Heusipp G, Greune L, Kim KS, Schmidt MA. Pertussis toxin transiently affects barrier integrity, organelle organization and transmigration of monocytes in a human brain microvascular endothelial cell barrier model. Cell Microbiol 2007;9(3):619–632.

59. Magira EE, Papaioakim M, Nachamkin I, Asbury AK, Li CY, Ho TW, Griffin JW, McKhann GM, Monos DS. Differential distribution of HLA-DQ beta/DR beta epitopes in the two forms of Guillain-Barre syndrome, acute motor axonal neuropathy and acute inflammatory demyelinating polyneuropathy (AIDP): identification of DQ beta epitopes associated with susceptibility to and protection from AIDP. J Immunol 2003;170(6):3074–3080.

60. Fekih-Mrissa N, Mrad M, Riahi A, Sayeh A, Zaouali J, Gritli N, Mrissa R. Association of HLA-DR/DQ polymorphisms with Guillain-Barre syndrome in Tunisian patients. Clinical neurology and neurosurgery 2014;121:19–22.

61. Jin PP, Sun LL, Ding BJ, Qin N, Zhou B, Xia F, Li L, Liu LJ, Liu XD, Zhao G, Wang W, Deng YC, Hou SX. Human Leukocyte Antigen DQB1 (HLA-DQB1) Polymorphisms and the Risk for Guillain-Barre Syndrome: A Systematic Review and Meta-Analysis. PloS one 2015;10(7):e0131374.

62. Hayat S, Jahan I, Das A, Hassan Z, Howlader ZH, Mahmud I, Deen Mohammad Q, Islam Z. Human leukocyte antigen-DQB1 polymorphisms and haplotype patterns in Guillain-Barre syndrome. Ann Clin Transl Neurol 2019;6(9):1849–1857.

