## Supplementary figures and images for "Development of a major histocompatibility complex class II conditional knockout mouse to study cell-specific and time-dependent adaptive immune responses in peripheral nerves"

### Supplemental Figure 1

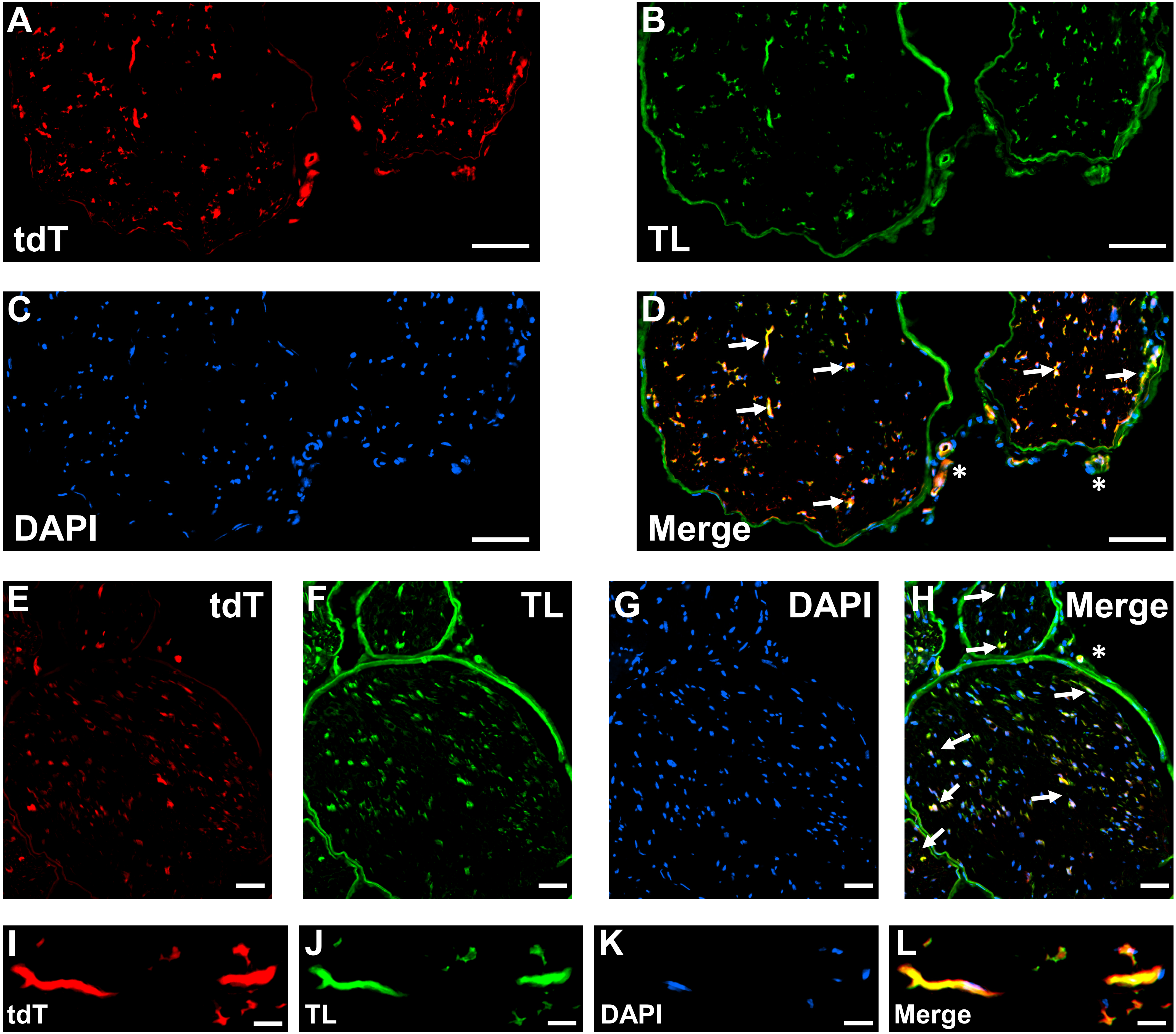

### Supplemental Figure 2

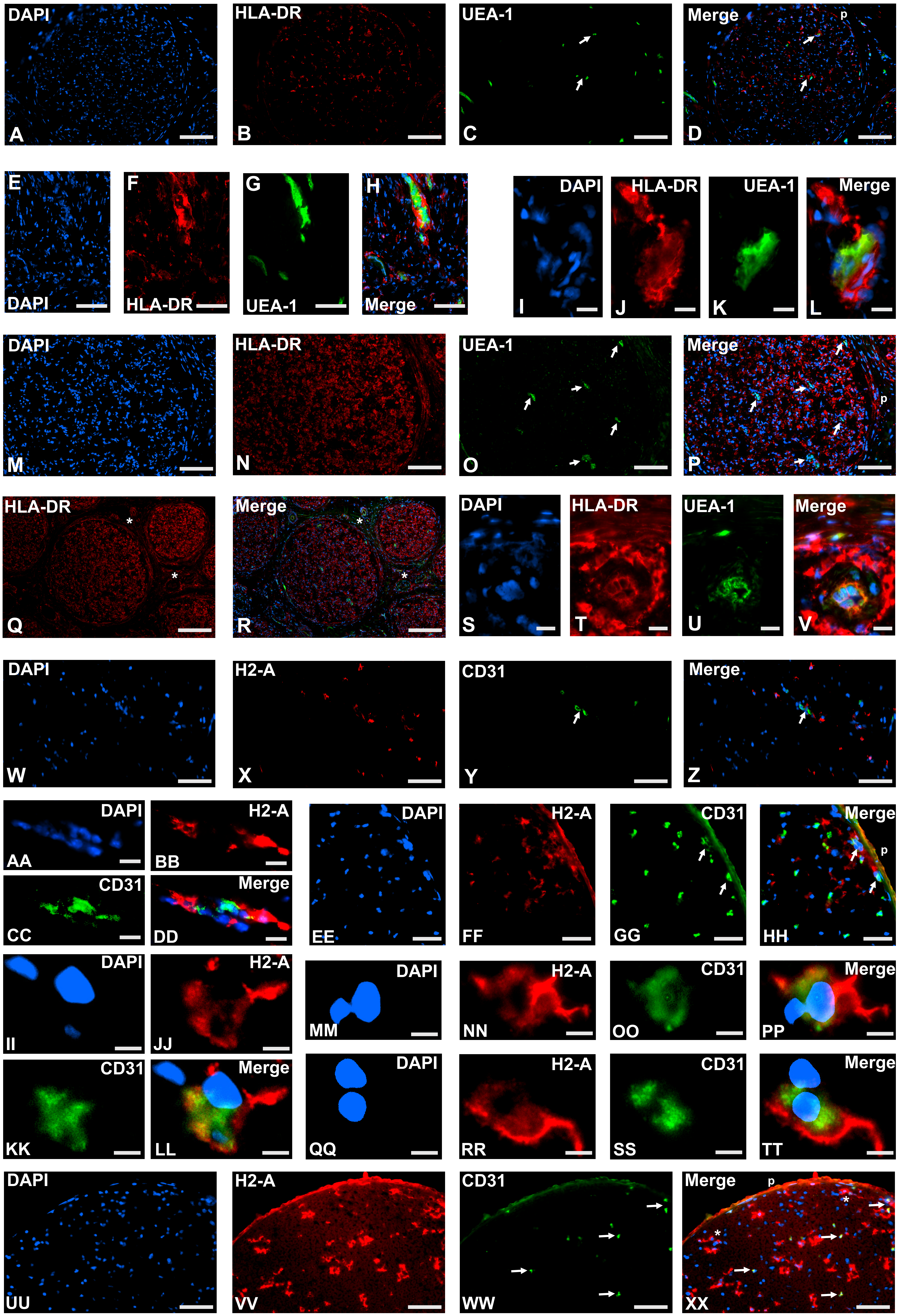

### Supplemental Figure 3

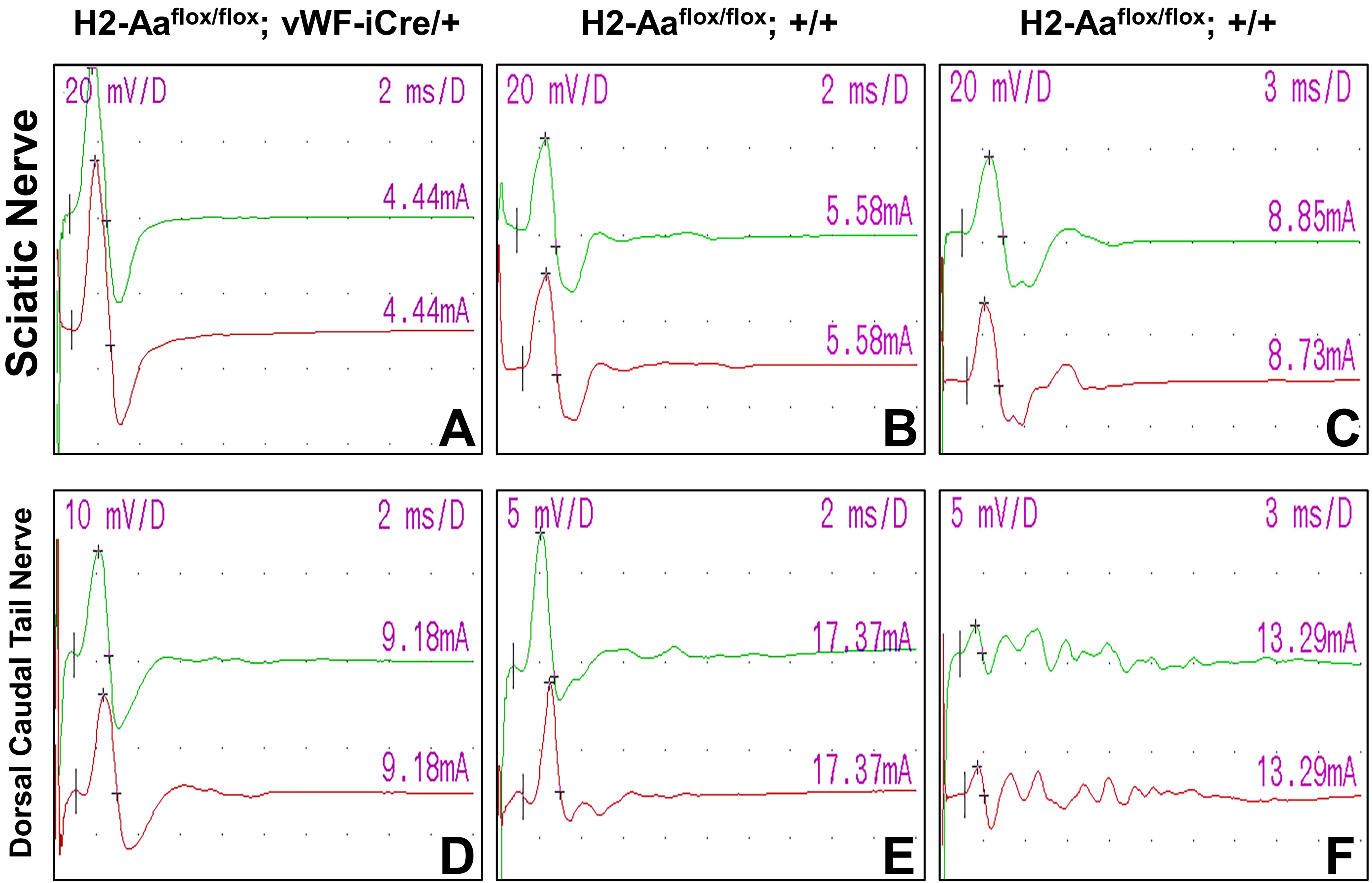

### Supplemental Figure 4

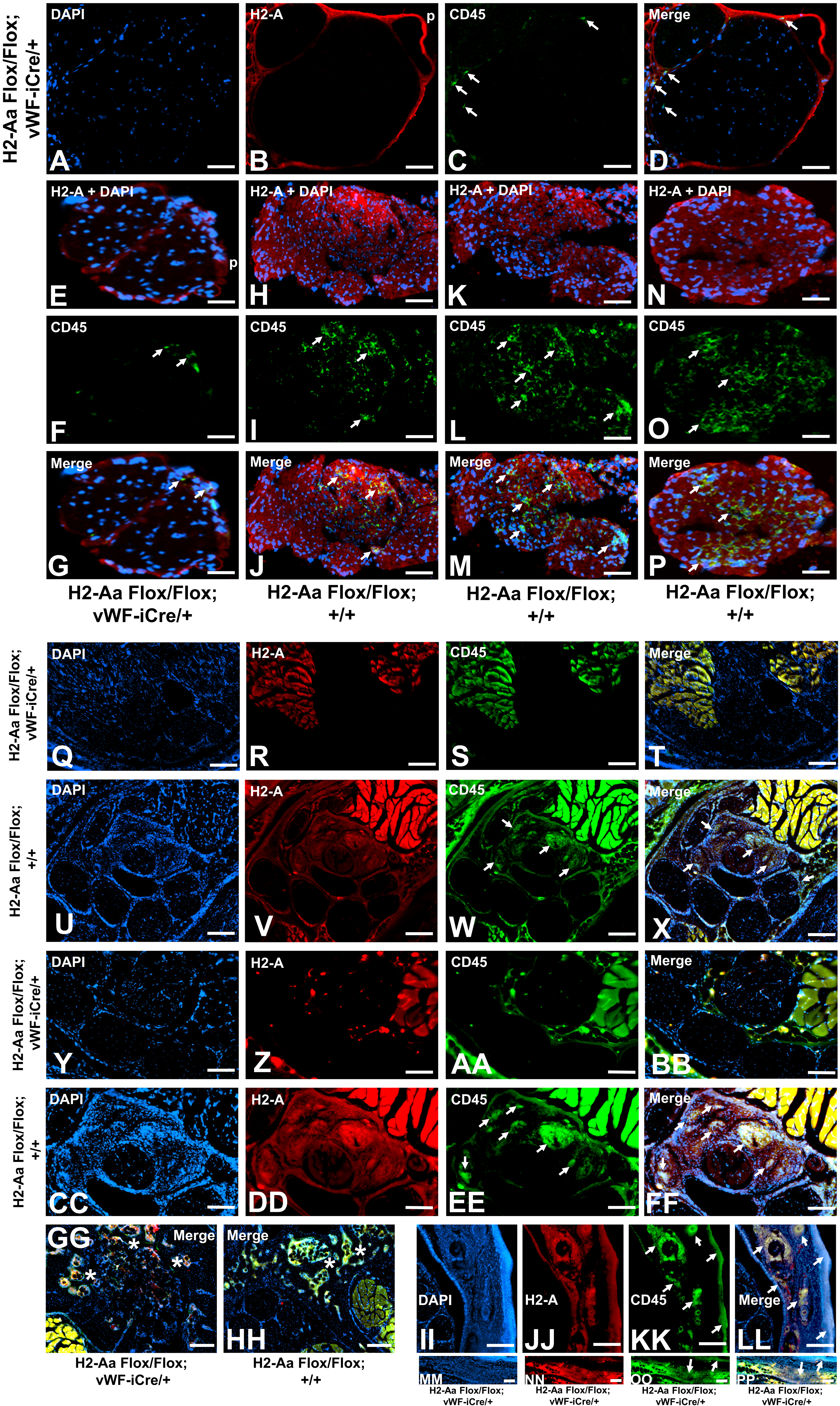
