## Supplemental Table 1 for "Development of a major histocompatibility complex class II conditional knockout mouse to study cell-specific and time-dependent adaptive immune responses in peripheral nerves"

| **PCR Process** | **TRANSGENE** | | |
| --- | --- | --- | --- |
|  | **Flpo recombinase** | **H2-Aa (SJL) 3’ and 5’ loxP sequences** | **vWF-iCre recombinase** |
| **Initiation/ Melting** | 95^o^C, 3 minutes | 94^o^C, 5 minutes | 95^o^C, 5 minutes |
| **Denaturation** | 95^o^C, 1 minute | 95^o^C, 30 seconds | 95^o^C, 30 seconds |
| **Annealing** | 56^o^C, 30 seconds | 55^o^C, 30 seconds | 55^o^C, 30 seconds |
| **Elongation** | 72^o^C, 1 minute | 72^o^C, 30 seconds | 72^o^C, 30 seconds |
| **Number of cycles** | 35 | 40 | 35 |
| **Amplification** | 72^o^C, 1 minute | 72^o^C, 5 minutes | 72^o^C, 5 minutes |

**Supplementary Table 1: PCR assay protocols**
