## Supplemental Table 2 for "Development of a major histocompatibility complex class II conditional knockout mouse to study cell-specific and time-dependent adaptive immune responses in peripheral nerves"

**Supplementary Table 2: Immunohistochemistry antibodies and lectins**

| **Target** | **Antibody/ Lectin** | **Clone** | **Concentration** | **Vendor** | **Catalog Number** |
| --- | --- | --- | --- | --- | --- |
| Human HLA-DR | Mouse IgG2b | LN3 | 2 µg/mL | ThermoFisher Scientific | MA5-11966 |
| Mouse MHC Class II H-2I-Ak/s/r | Rat IgG2b | ER-TR2 | 10 µg/mL | Bio-Rad Laboratories | MCA2400 |
| Human endothelial cell marker  (binds to α-fucose) | UEA-1 FITC | - | 10 µg/mL | ThermoFisher Scientific | L32476 |
| Mouse endothelial cell marker  (binds to [GlcNAc] 1,3-N-acetylglucosamine) | TL-DyLight 488 | - | 5 µg/mL | ThermoFisher Scientific | L32470 |
| Mouse CD31 | Rabbit polyclonal IgG | - | 2.6 µg/mL | ThermoFisher Scientific | PA5-16301 |
| Mouse CD45 | Rabbit polyclonal IgG | - | 6.7 µg/mL | Abcam | ab10558 |
| Mouse IgG | Goat polyclonal IgG AlexaFluor^TM^ 594 | - | 4 µg/mL | ThermoFisher Scientific | A32742 |
| Rat IgG | Goat polyclonal IgG AlexaFluor^TM^ 594 | - | 4 µg/mL | ThermoFisher Scientific | A48264 |
| Rabbit IgG | Goat polyclonal IgG AlexaFluor^TM^ 488 | - | 4 µg/mL | ThermoFisher Scientific | A-11034 |
